# Adolescent intermittent ethanol exacerbates Aβ with age in the dorsal hippocampus of female TgF344-AD rats

**DOI:** 10.1101/2025.10.20.683498

**Authors:** Stephen M. Day, Nicole L. Reitz, Lisa M. Savage

**Affiliations:** Savage Department of Psychology, Binghamton University, State University of New York, New York, USA

**Keywords:** ethanol, adolescence, amyloid-β, cortex, hippocampus, Alzheimer’s disease

## Abstract

**Background:** Alcohol misuse increases Alzheimer’s disease (ΑD) risk, however the mechanisms linking these conditions are unknown. In rodents, chronic and acute ethanol increases amyloid-β (Aβ), however those studies have been limited to a single sex or brain region.

**Objective:** This study explored how adolescent intermittent ethanol (AIE), alters Aβ in multiple regions of the brain in female and male TgF344-AD rats as they age.

**Methods:** From P28-P58, female and male TgF344-AD rats were administered either water (CON) or 5.0 g/kg ethanol (AIE; 20% ethanol w/v) via intragastric gavage on a 2-day on/off cycle. In Experiment 1, Aβ was measured in the medial prefrontal cortex (mPFC), orbitofrontal cortex (OFC), piriform cortex (PC), entorhinal cortex (EC), ventral hippocampus (vHPC), and dorsal hippocampus (dHPC) in 6- and 10-month-old rats. In Experiment 2, in vivo microdialysis was used in 3-month-old female rats to measure how ethanol directly modulates Aβ levels in the dHPC.

**Results:** In the OFC, PC, EC, vHPC, and dHPC, Aβ40 and Aβ42 was higher in 6-month-old female TgF344-AD rats compared to males. However, at 10 months Aβ40 and Aβ42 levels were only elevated in the dHPC of AIE-treated females, compared to all other groups. An acute ethanol challenge at 3 months selectively evoked a sustained increase in ISF Aβ40 levels in AIE-treated females.

**Conclusions:** In aged females, the dHPC is a region sensitive to ethanol-associated Aβ pathology. This may be due to disruptions in Aβ clearance in early life, which may have an additive effect on Aβ aggregation over the lifespan.

## Introduction

Alzheimer’s disease (AD) is the most common form of dementia, affecting more than 7.2 million adults in the United States alone [1]. AD-related pathology is characterized by the accumulation of amyloid-β (Aβ) into extracellular amyloid plaques, the accumulation of intracellular neurofibrillary tangles, and neurodegeneration [2,3]. These pathological events begin to accrue ∼15 years before the onset of clinical diagnosis [4], thus it is important to identify modifiable risk factors that accelerate the onset of clinical symptoms and cognitive decline associated with AD. Epidemiological studies have identified alcohol misuse as a risk factor for AD and dementia [5–10]. Conversely, a few studies suggest that low-to moderate drinking may not impact AD risk at all [11,12]. Despite this, there is a growing body of preclinical evidence demonstrating that chronic heavy ethanol exposure promotes AD-related pathology in rodent models of AD-like pathology [13–15]. Thus, it is important to identify how ethanol misuse contributes to AD-related pathology throughout all stages of the lifespan.

Binge-like ethanol exposure is characterized by excessive alcohol intake followed by periods of abstinence and is a risk factor for alcohol misuse during adulthood. Adolescence is an especially vulnerable period of neurodevelopment and binge-like alcohol consumption during this period can have long-lasting consequences on brain health and structure [16,17]. In preclinical studies binge-like ethanol exposure, increases Aβ protein levels in the hippocampus with advancing age [18,19]. While informative, these studies are limited in that they have only explored how adolescent binge-like ethanol impacts Aβ levels in females and in a single region of the brain. Despite a growing body of research on alcohol–AD interactions, the role of sex remains critically understudied.

Few investigations have systematically examined sex as a biological variable in preclinical models, representing a significant gap in the field. This omission is particularly consequential given robust evidence that women exhibit a higher incidence of AD and demonstrate more rapid trajectories of cognitive decline and neurodegeneration following diagnosis [20–22]. Addressing this gap is essential for advancing mechanistic understanding and developing sex-specific preventive and therapeutic strategies.

In this study, we built on that previous work by investigating how adolescent intermittent ethanol (AIE), a model of binge-like ethanol consumption, impacts Aβ levels across multiple regions of the brain and across the lifespan [19,23,24]. The model chosen was the TgF344-AD rat model [25–28], as it is a robust platform for investigating alcohol– AD interactions because it recapitulates multiple hallmark pathologies of AD, including age-related progressive amyloid-β deposition, tau hyperphosphorylation, neuroinflammation, and cognitive decline. TgF344-AD rats exhibit both amyloid and tau pathologies in a temporal sequence that more closely parallels human disease progression, providing enhanced translational relevance [25,29,30]. In this model, there is a progressive pattern of AD-related pathology and abnormal activity across the hippocampus and cortical regions [31].

Specifically, in this series of studies, we examined changes in Aβ across the orbitofrontal cortex (OFC), medial prefrontal cortex (mPFC), piriform cortex (PC), entorhinal cortex (EC), ventral hippocampus (vHPC), and dorsal hippocampus (dHPC) in both female and male TgF344-AD rats. We found that 6-month-old female TgF344-AD rats showed higher levels of Aβ40, Aβ42, and/or Aβ42/40 across most regions investigated, except for the mPFC. By 10 months of age, there were greater Aβ40 and Aβ42 levels, but only in the dHPC did AIE-treated females display exaggerated Aβ40 and Aβ42 levels. To examine ethanol sensitivity during the prodromal period, in vivo microdialysis was used to explore how ethanol directly influences interstitial fluid (ISF) Aβ levels in female TgF344-AD rats with a history of binge-like ethanol. Here, AIE-treated 3-month-old female rats showed a prolonged, dose-dependent increase in Aβ40 after an acute ethanol challenge. This suggests that AIE may disrupt Aβ clearance mechanisms, and potentially exacerbating Aβ deposition later in life. These studies build on previous work by showing sex-dependent increases in Aβ during the prodromal period of AD pathogenesis across multiple regions of the brain. Importantly, these studies show sex- and AIE-dependent increases in Aβ40 and Aβ42 in the dHPC of middle-aged animals. This study distinguishes the dHPC from the vHPC as a particularly vulnerable region in females to AIE-associated increases in Aβ. Lastly, these studies suggest that a history of ethanol misuse during adolescence may have long-lasting effects on Aβ production or clearance mechanisms in the brain that may be exacerbated with each additional exposure during adulthood.

## Methods

### Subjects

Male (n = 41) and female (n = 70; Experiment 1 = 38; Experiment 2 = 42) TgF344-AD rats were bred in-house at Binghamton University from F344 dams (Charles River, Kingston, NY) and TgF344-AD males (RRRC, Colombia, MO). No more than 1 rat per sex per litter was assigned to an experimental treatment condition. At postnatal day 21 (P21), pups were weaned and genotyped using ear-punched tissue sent to Transnetyx (Cordova, TN) for genotyping of APPswe and PSEN1ΔE9. Rats were housed with one or two other same-sex conspecifics in a humidity and temperature-controlled colony room set to 20°C on a 12-hour light/dark cycle. All experimental procedures were approved by the Institutional Animal Care and Use Committee (IACUC) at the State University of New York at Binghamton and followed the National Institutes of Health (NIH) Guide for Care and Use of Laboratory Animals.

### Adolescent intermittent ethanol (AIE)

Rats were randomly assigned to control (CON) or AIE treatment groups. From on P28 to P58, male and female rats were administered either water (CON) or 20% ethanol (AIE; w/v, in tap water) via intragastric gavage (i.g.) at a dose of 5.0 g/kg on a 2-day on/off cycle (Figure 2a), as previously described [32,33]. One hour after the 8th gavage, ∼50-100 μL of blood was taken from the saphenous vein and blood ethanol concentrations (BECs) were measured with an AM1 Alcohol Analyzer (Analox Instruments, Stourbridge UK). In experiment 1, rats were aged to 6- or 10-months. In experiment 2, 32 female rats underwent cannula implantation surgery at 3-months and were used for in vivo microdialysis experiments (see below).

### Experiment 1: Soluble Aβ extraction

Soluble Aβ (sAβ) fraction was extracted as described previously [34]. Briefly, 6- and 10-months rats were humanely euthanized via rapid decapitation, and brains were immediately removed and flash-frozen then stored at -80°C. Tissue punches were collected via cryostat (Lecia Biosystems, Deer Park, Il), from the mPFC, OFC, PC, EC, vHPC, and dHPC. These regions were selected based on Paxinos & Watson Stereotaxic Coordinates (Supplementary Figure 1) [35]. Tissue was weighed and 20 μL/mg of a DPBS buffer containing Complete protease inhibitor (Roche, Indianapolis, IN) and phosStop phosphatase inhibitor (Roche) was added. The samples were homogenized with a bead homogenizer for 50 seconds and centrifuged at 15,000 rpm for 30 minutes. The supernatant was removed, and total protein concentrations were analyzed using a Pierce BCA protein assay kit (Thermo Scientific, Waltham, MA; P5934129). The remaining lysate was used for Aβ40 and Aβ42 ELISA (see below).

### Experiment 2: In vivo microdialysis

We previously showed that ethanol-naïve APP/PS1 male mice have a transient increase in Aβ40 levels after an acute ethanol exposure [13]. Here, given our sex-dependent findings in Experiment 1, only female 3-month-old TgF344-AD rats were used to measure how acute ethanol alters Aβ40 levels prior to the onset of brain Aβ deposition. Interstitial fluid (ISF) was continuously collected from the dHPC before and after ethanol exposure using in vivo microdialysis as previously described [13,36]. Approximately five days prior to acute ethanol exposure, guide cannulas (AG-12; Amuza, San Diego, CA) were stereotaxically implanted into the dHPC (from bregma, A/P: 3.8 mm; M/L: 3.43 mm; D/V: 2.77 mm; at 12⁰ angle) and secured into place with dental cement. 12 hours prior to ethanol, rats were transferred to sampling cages (Bioanalytical Systems, West Lafayette, IN), and microdialysis probes (FZ-12-02; 50 kDa molecular weight cut off; Amuza) were inserted into the guide cannula, connected to a syringe pump and infused with 0.15% bovine serum albumin (BSA, Sigma-Aldrich, St. Louis, MO) in artificial cerebrospinal fluid (aCSF; 1.3 mM CaCl2, 1.2 mM MgSO4, 3 mM KCl, 0.4 mM KH2PO4, 25 mM NaHCO3 and 122 mM NaCl; pH = 7.35) at a flow rate of 1 μL/min. Hippocampal ISF was collected every 1.5 hours, beginning at 8:00 pm. 12.5 hours later, mice were administered 0.9% saline or 2.0 g/kg ethanol (i.p) from a 15% ethanol (w/v; in 0.9% saline) stock solution.

Collected ISF for 9 hours was stored at 20°C for Aβ40 ELISA (see below). Rats were then returned to their home cages and were humanely euthanized via rapid decapitation a week later.

### Aβ40 and Aβ42 ELISAs

Brain lysate and ISF samples from were analyzed for Aβ40 and/or Aβ42 using sandwich ELISAs as previously described [13,36–38]. Briefly, Aβ40 and Aβ42 were quantified using monoclonal capture antibodies (generous gifts from Drs. Hong Jiang and David Holtzman at Washington University in St. Louis, MO) targeted against amino acids 33-40 (HJ2, Aβ40) or amino acids 35-42 (HJ7.4, Aβ42). For detection, a biotinylated monoclonal antibody against the central domain amino acids 13-28 (HJ5.1B) was used, followed by streptavidin-poly-HRP40. Assays were developed using Super Slow TMB (Sigma-Aldrich). Plates were read on a Tecan Infinite M200 Pro plate reader at 650 nm. Aβ concentrations from brain lysate was normalized to total protein (pg/μg) while ISF Aβ concentrations were normalized to baseline Aβ levels (% baseline).

### Statistical Analysis

Statistical analyses were performed with GraphPad Prism 10 (GraphPad Software Inc, San Diego CA) or SPSS (IBM, Armonk NY). Two-way ANOVAs were used to analyze the differences between subject factors (Sex [Male, Female] and Treatment [CON, AIE]) and post-hoc analyses (Fishers LSD) were performed for assessing specific group comparisons. 3-way repeated measures ANOVAs were used to analyze body weight (Gavage number, Treatment, and Sex) and ISF Aβ40 levels (Treatment, Time, and EtOH/Saline challenge). Grubbs outlier tests were performed on all data (α = 0.05) and outliers were excluded from analyses. The level of statistical significance weas set at p ≤ 0.05. Data are expressed as means ±SEM. Complete statistical results can be found in Supplementary Table 1.

## Results

### Body weight and blood ethanol content (BEC)

Body weights were recorded before each gavage session (Figure 1b). As expected, body weights in males increased at a higher rate than females. While there were no differences between control or AIE-treated females throughout the gavage period, post-hoc tests showed that control males had higher body weights than AIE-treated males after the 9th gavage (Time x Treatment x Sex: p < 0.0001; F(1,15) = 3.476). This is consistent with our lab’s previous reports [33]. One hour after the 8^th^ gavage, we collected blood from the saphenous vein and measured blood ethanol concentrations (BECs) from AIE-treated rats and no sec differences in BECs were found (Figure 1c).

**Figure 1:**
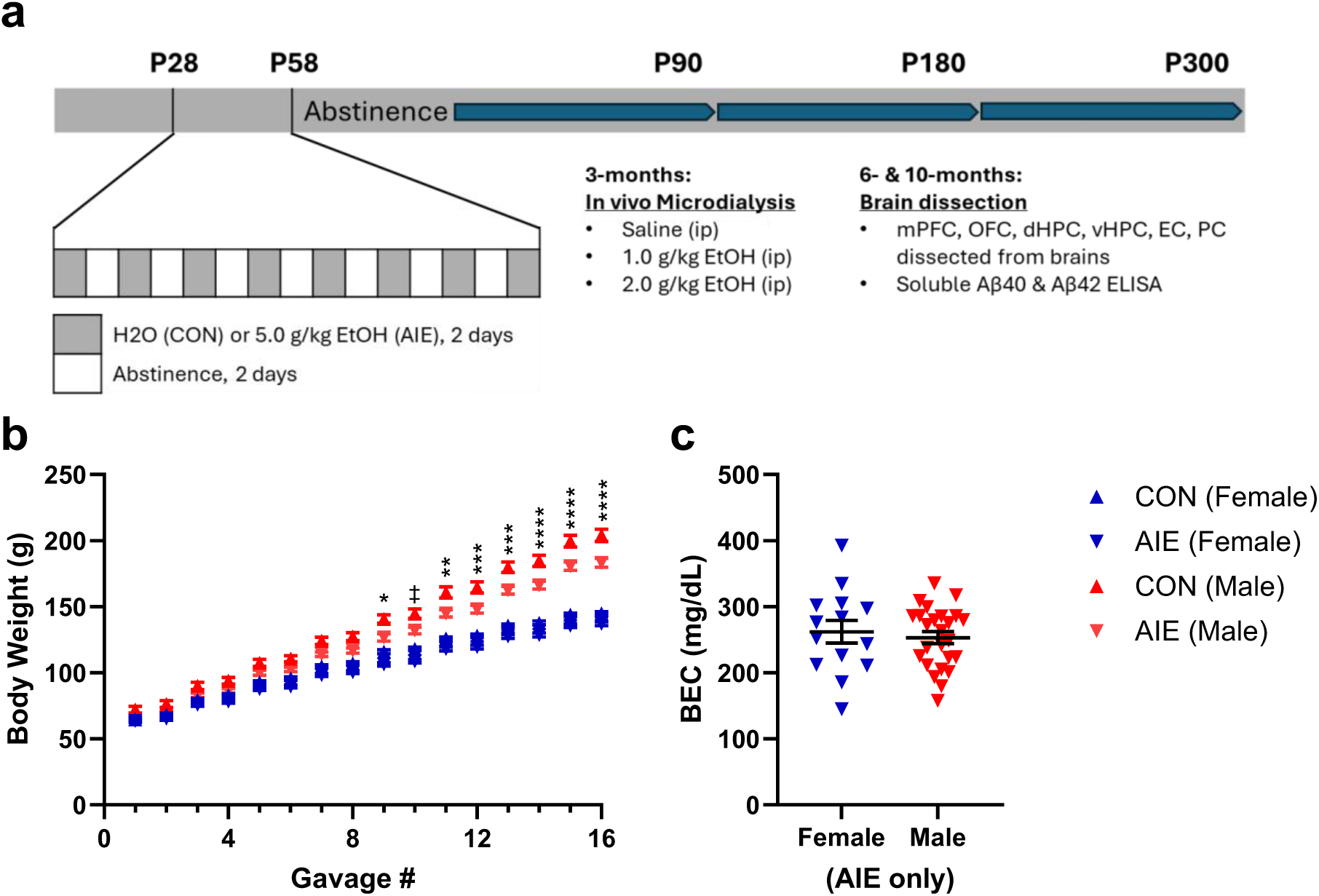
Body weight and BEC. a) Timeline for experimental protocol. Full details are provided in the methods. b) Mean body weights for CON- and AIE-treated female and male TgF344-AD rats. Male rats showed greater body weights with CON males having greater body weights than AIE males (3-way ANOVA, Main effect of time: p < 0.0001; Time x treatment interaction: p < 0.0001; time x sex interaction: p < 0.0001; time x treatment x sex interaction: p < 0.0001). c) There were no differences in BECs between AIE-treated females and males (Unpaired t-test, p = 0.6500 (t=0.4605, df=20.53)). ‡p < 0.10, *p < 0.05, ** p < 0.01, ***p < 0.001, ****p < 0.0001

**Figure 2:**
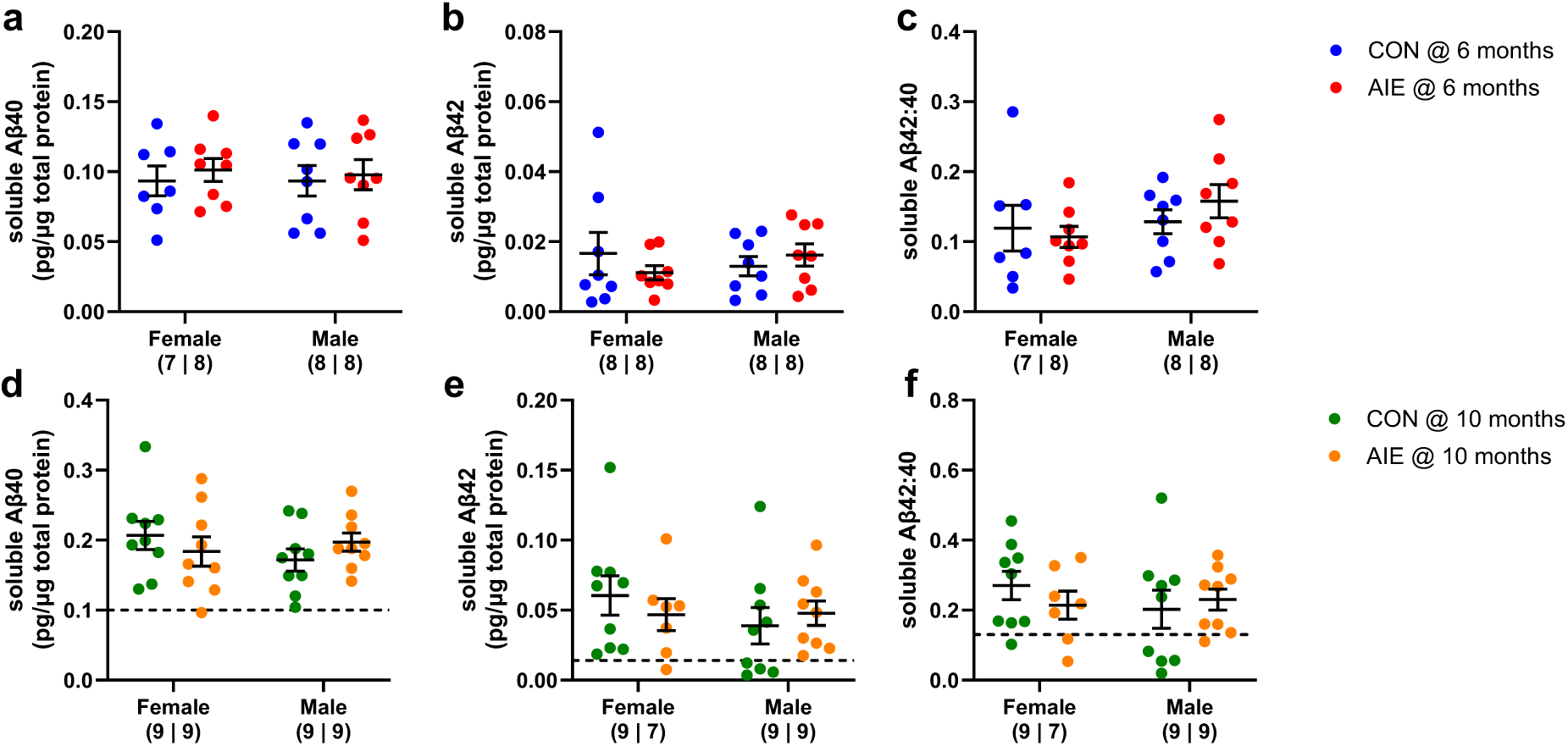
In the medial prefrontal cortex (mPFC), neither AIE-treatment nor sex had no impact on Aβ levels in 6- or 10-month-old TgF344-AD rats. In 6-month-old TgF344-AD rats there were no differences in Aβ40, Aβ42, or Aβ42/40 between groups (a-c). In 10-month-old TgF344-AD rats (d-f) there were no differences in Aβ40, Aβ42, or Aβ42/40 between groups. There was an age-dependent increase in Aβ40 levels (3-way ANOVA, Main effect of age: p < 0.0001). Dashed line represents mean values of 6-month-old rats.

### Experiment 1: Sex-, age-, and regional differences in Aβ levels of control- and AIE-treated TgF344-AD rats

In the mPFC, there were no differences in Aβ40 (Figure 2a) or Aβ42 levels (Figure 2b) as a function of treatment or sex in 6-month-old rats. There were also no differences between groups in the Aβ42/40 ratio in 6-month-old animals (Figure 2c). Similarly, in 10-month-old rats, there were also no differences between groups in Aβ40 (Figure 2d) or Aβ42 (Figure 2e) levels, nor were there any differences in the Aβ42/40 ratio (Figure 2f) in the mPFC. However, Aβ40 levels did increase with age (3-way ANOVA: Main effect of age, F(1,60) = 59.3292, p < 0.0001) and Aβ42 levels showed a trend towards age-dependent increases (3-way ANOVA: Main effect of Age, F(1,60) = 3.4624, p = 0.0677). Thus, in the mPFC, progression of AD-related pathology is not dependent upon either sex or influenced by a history of AIE.

We next examined Aβ changes in the OFC. Although at 6-months there were no Sex or Treatment differences in Aβ40 levels (Figure 3a), TgF344-AD female rats to have higher Aβ42 levels compared to AD males (Figure 3b; Main effect of Sex: F(1,28) = 5.127, p = 0.0315). There was also a trend towards a main effect of Sex in the Aβ42/40 ratio in the OFC, with females having a higher ratio compared to males (Figure 3c; Main effect of Sex: F (1, 28) = 3.746; p = 0.0631). In 10-month-old TgF344-AD rats, there were no Sex or Treatment differences in Aβ40, or Aβ42 levels (Figure 3d-f). However, there was a trend towards a Treatment x Sex interaction in the Aβ42/40 ratio (F (1, 29) = 3.234; p = 0.0825). Control-treated females had higher Aβ42/40 compared to AIE-treated females (p = 0.0971), and control-treated males (p = 0.0555). In the OFC, there was also an age-dependent increase in Aβ40 (Main effect of ageF(1,59) = 17.8284, p = 0.0001), Aβ42 (Main effect of Age: F(1,59) = 21.3660, p < 0.0001), and the Aβ42/40 ratio (Main effect of Age: F(1,59) = 46.2188, p < 0.0001).

**Figure 3:**
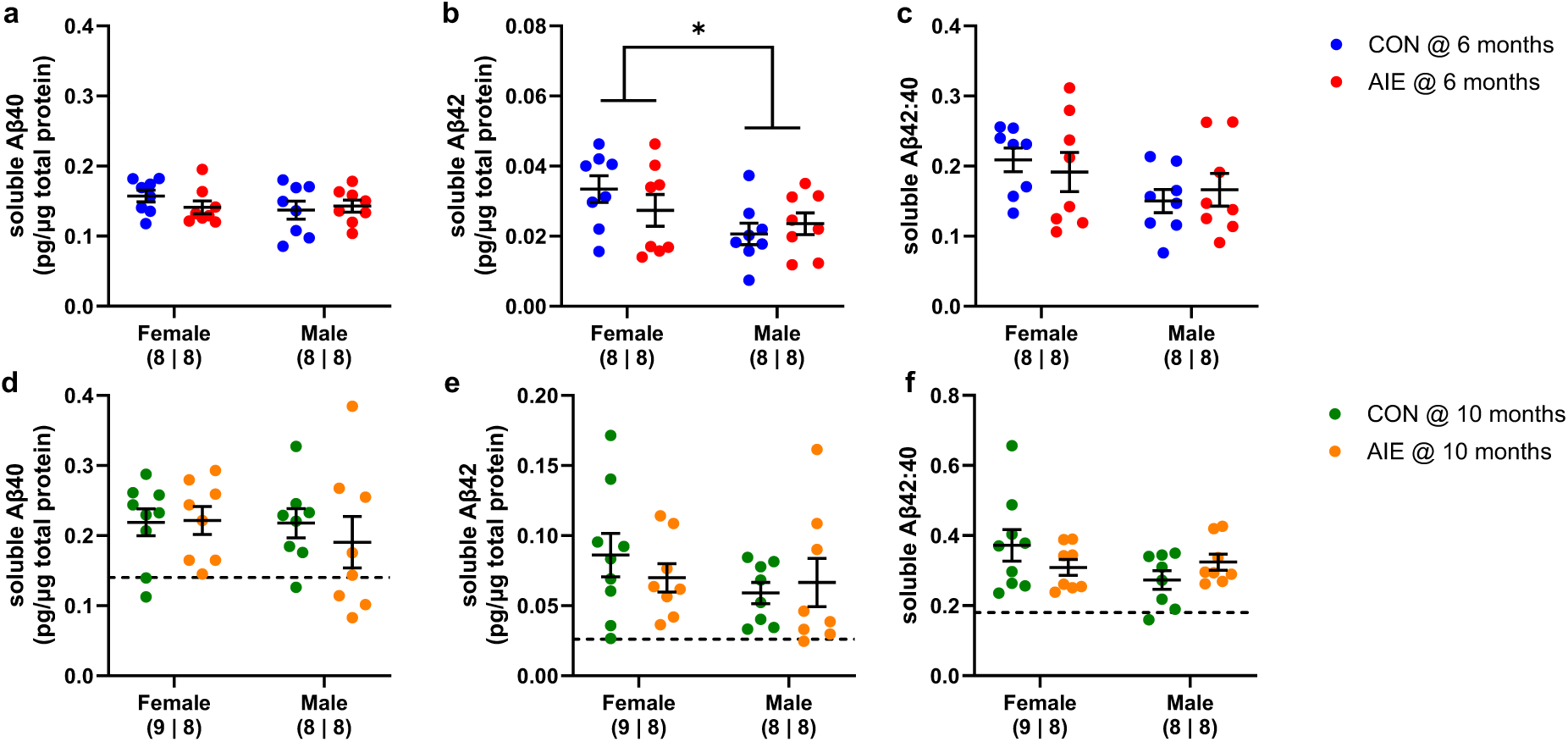
In the orbitofrontal cortex (OFC), 6-month-old female TgF344-AD rats showed higher soluble Aβ42 levels than males. In 6-month-old TgF344-AD rats there were no differences in soluble Aβ levels between groups (a). However female rats showed higher Aβ42 levels than males (b; 2-way ANOVA, Main effect of sex: p = 0.0315); and there was a trend towards increased Aβ42/40, favoring females over males (c; 2-way ANOVA, Main effect of sex: p = 0.0631). In 10-month-old TgF344-AD rats there were no differences in Aβ40 or Aβ42 (d-e), but there was a trend towards a sex x treatment interaction with the Aβ42/40 ratio (f; p = 0.0825). There were age-dependent increases in Aβ40 (a,d; 3-way ANOVA, Main effect of age: p = 0.0001), Aβ42 (b,e; 3-way ANOVA, Main effect of age: p < 0.0001), and Aβ42/40 (c,f; 3-way ANOVA, Main effect of age: p < 0.0001). Dashed line represents mean values of 6-month-old rats. *p < 0.05

In the PC, 6-month-old AD females showed higher Aβ40 (Main effect of Sex: F(1,28) = 8.687; p = 0.0064) and Aβ42 (Main effect of Sex: F(1,24) = 9.111; p = 0.0059) levels and a greater Aβ42/40 ratio (Main effect of sex: F(1,23) = 9.583; p = 0.0051) than TgF344-AD males (Figure 4a-c). Post-hoc tests also showed that AIE-treated TgF344-AD females had a higher Aβ42/40 ratio than control TgF344-AD males (p = 0.0377). In 10-month-old TgF344-AD rats, there were no Treatment or Sex differences in Aβ40 or Aβ42 levels, or in the Aβ42/40 ratio (Figure 4d-f). Interestingly, Aβ40 and Aβ42 levels, as well as the Aβ42/40 ratio more than doubled from 6- to 10-months in all animals. Here, a 3-way ANOVA revealed an age-dependent increase in Aβ40 (Main effect of age: F(1,56) = 166.8987, p < 0.0001), Aβ42 (Main effect of Age: p < 0.0001; F(1,54) = 344.3122), and Aβ42/40 (Main effect of Age: F(1,54) = 654.2878, p < 0.0001). While early sex-related differences were no longer detectible, PC is a particularly vulnerable to age-related increases in AD-related pathology in TgF344-AD rats. This is consistent with previous studies showing age-dependent increases in PC plaque deposition of Tg2576 mice [37].

**Figure 4:**
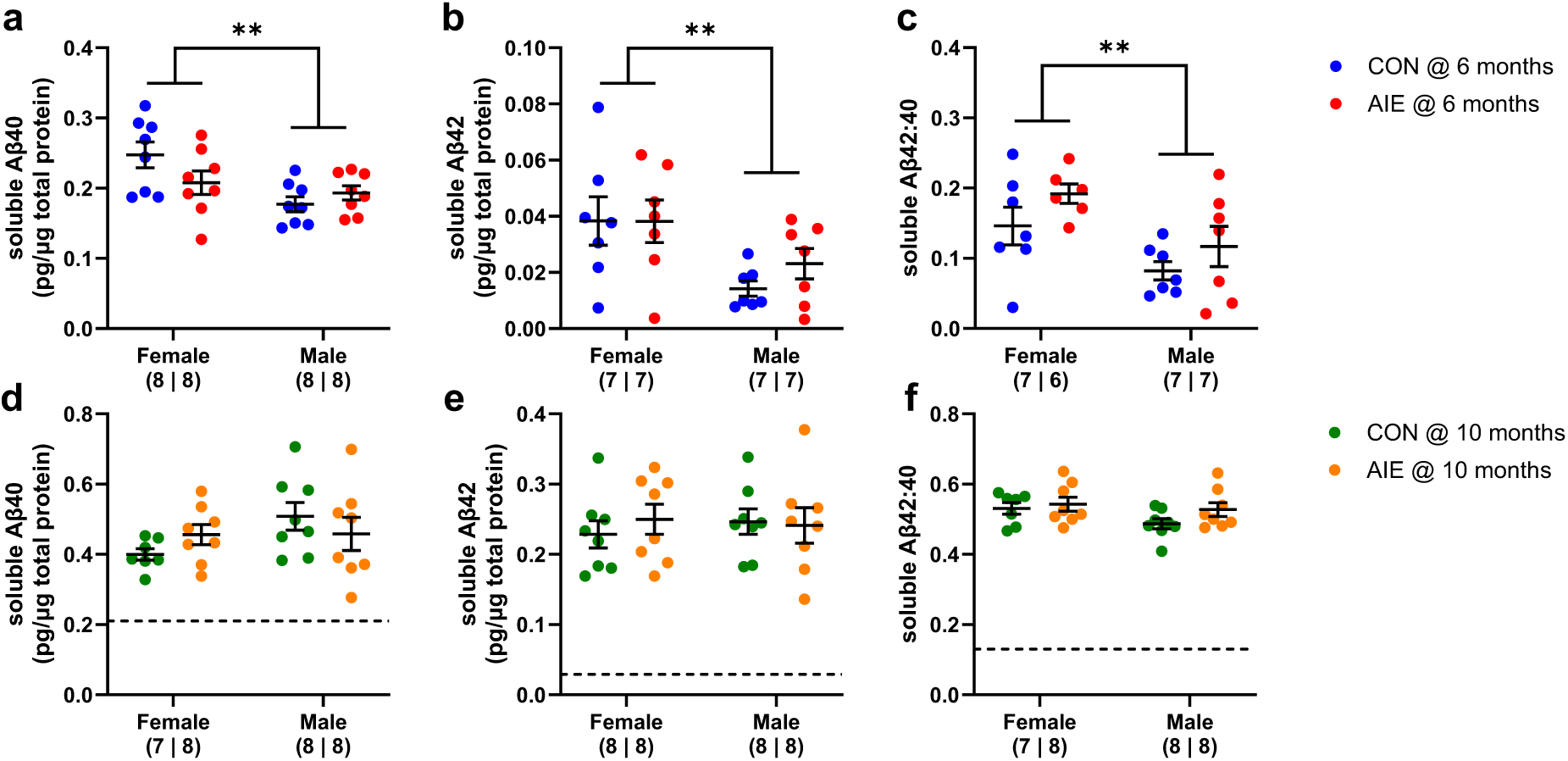
In the piriform cortex (PC), 6-month-old female TgF344-AD rats showed higher soluble Aβ40, Aβ42, and Aβ42/40 than males. In 6-month-old TgF344-AD female rats had higher Aβ40 levels (a; 2-way ANOVA, Main effect of sex: p = 0.0064), Aβ42 (b; 2-way ANOVA, Main effect of sex: p = 0.0059), and a higher Aβ42/40 ratio than males (c; 2-way ANOVA, Main effect of sex: p = 0.0051). In 10-month-old TgF344-AD rats (d-f) there were no differences in Aβ40, Aβ42, or Aβ42/40 between groups. There were also age-dependent increases in Aβ40 (3-way ANOVA, Main effect of age: p = 0.0001), Aβ42 (3-way ANOVA, Main effect of age: p < 0.0001), and Aβ42/42 (Main effect of age: p < 0.0001). Dashed line represents mean values of 6-month-old rats. **p < 0.01; *p < 0.05

In the EC, 6-month-old female rats showed greater Aβ40 levels compared to males (F (1, 28) = 10.20, p = 0.0035; Figure 5a). Six month-old females also showed greater EC Aβ42 levels compared to males (Main effect of Sex: F(1,28) = 8.006; p = 0.0085; Treatment x Sex interaction: F (1, 28) = 4.872, p = 0.0357; Figure 5b); Post-hoc tests revealed that control females had higher Aβ42 levels than control males (p = 0.0069) in the EC. Meanwhile there were no differences in the Aβ42/40 EC ratio in 6-month-old rats. In contrast, in 10-month-old rats, there were no differences in EC Aβ40 or Aβ42 levels, nor were there any differences in the Aβ42/40 ratio between groups (Figure 5d-f). Again, early sex-related differences in EC Aβ40 and Aβ42 levels were no longer present in 10-month-old animals. A 3-way ANOVA revealed an age-dependent increase in Aβ40 (Main effect of age: F(1,60) = 103.9577, p = 0.0001), Aβ42 (Main effect of Age: F(1,59) = 241.8610, p < 0.0001;), and Aβ42/40 (Main effect of Age: F(1,59) = 235.8186, p < 0.0001).

**Figure 5:**
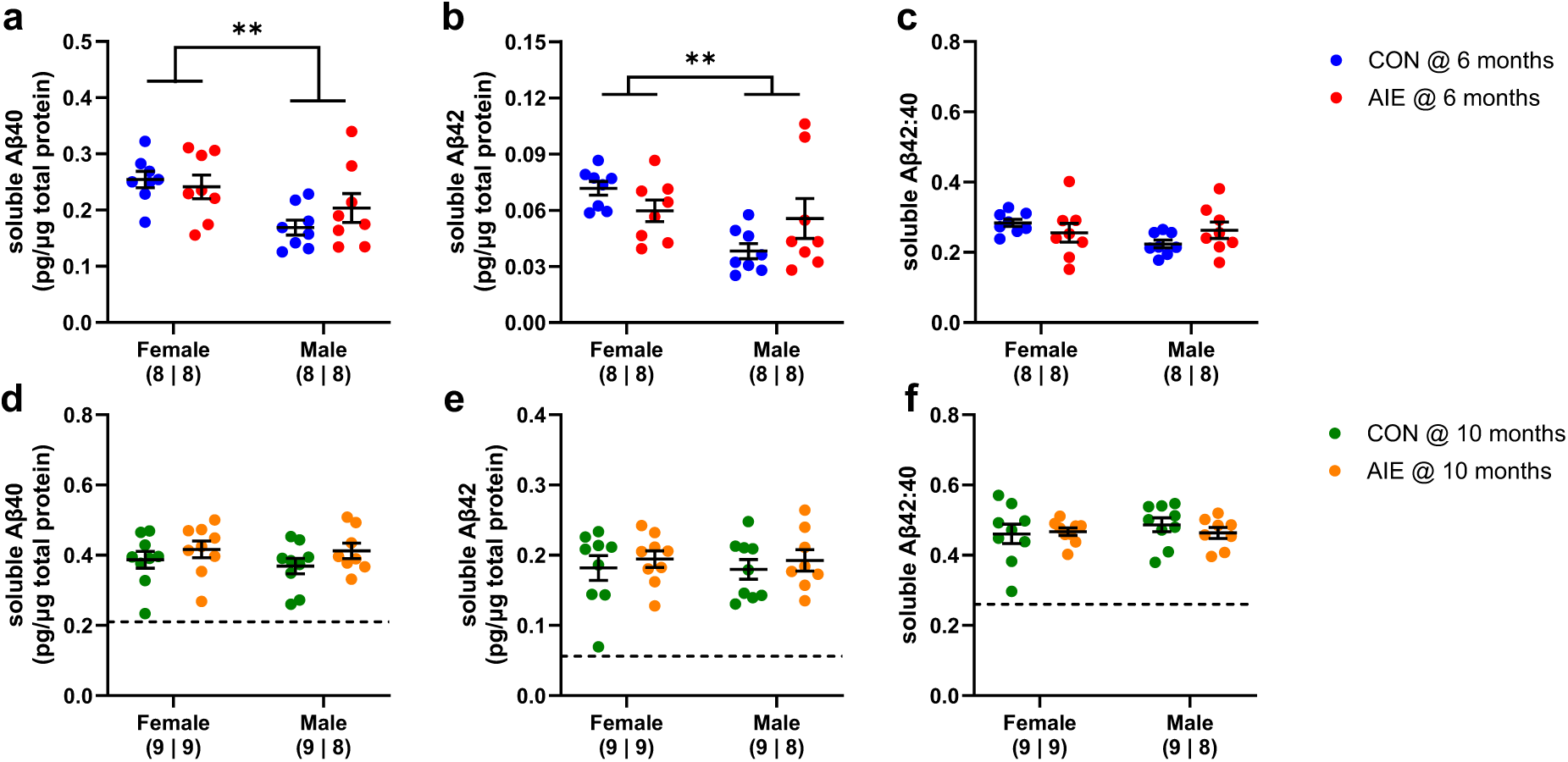
In the entorhinal cortex (EC), 6-month-old female TgF344-AD rats showed higher soluble Aβ40 and Aβ42 levels than males. In 6-month-old TgF344-AD rats females had higher Aβ40 (a; 2-way ANOVA, Main effect of sex: p = 0.0035) and Aβ42 levels than males (b; 2-way ANOVA, Main effect of sex: p = 0.0085; Treatment x sex interaction: p = 0.0357), and there was a trend towards a Treatment x Sex interaction with Aβ40/42 with CON-females having higher than CON-males (c; 2-way ANOVA, Treatment x sex interaction: p = 0.0933). In 10-month-old TgF344-AD rats there were no differences in Aβ40, Aβ42, or Aβ42/40 (d-f). There were age-dependent increases in Aβ40 (3-way ANOVA, Main effect of age: p = 0.0001), Aβ42 (3-way ANOVA, Main effect of age: p < 0.0001), and Aβ42/40 (3-way ANOVA, Main effect of age: p < 0.0001). Dashed line represents mean values of 6-month-old rats. ***p < 0.001; **p < 0.01

In the vHPC, 6-month-old TgF344-AD female rats showed greater Aβ40 (Figure 6a; Main effect of Sex: F(1,24) = 9.046; p = 0.0061) and Aβ42 levels (Figure 6b; Main effect of Sex: F(1,26) = 18.70; p = 0.0002) compared to AD males. TgF344-AD females also showed a higher Aβ42/40 ratio compared to males (Figure 6c; Main effect of Sex: F(1,25) = 16.10; p = 0.0005; Treatment x Sex interaction: F(1,54) = 6.338; p = 0.0186).

**Figure 6:**
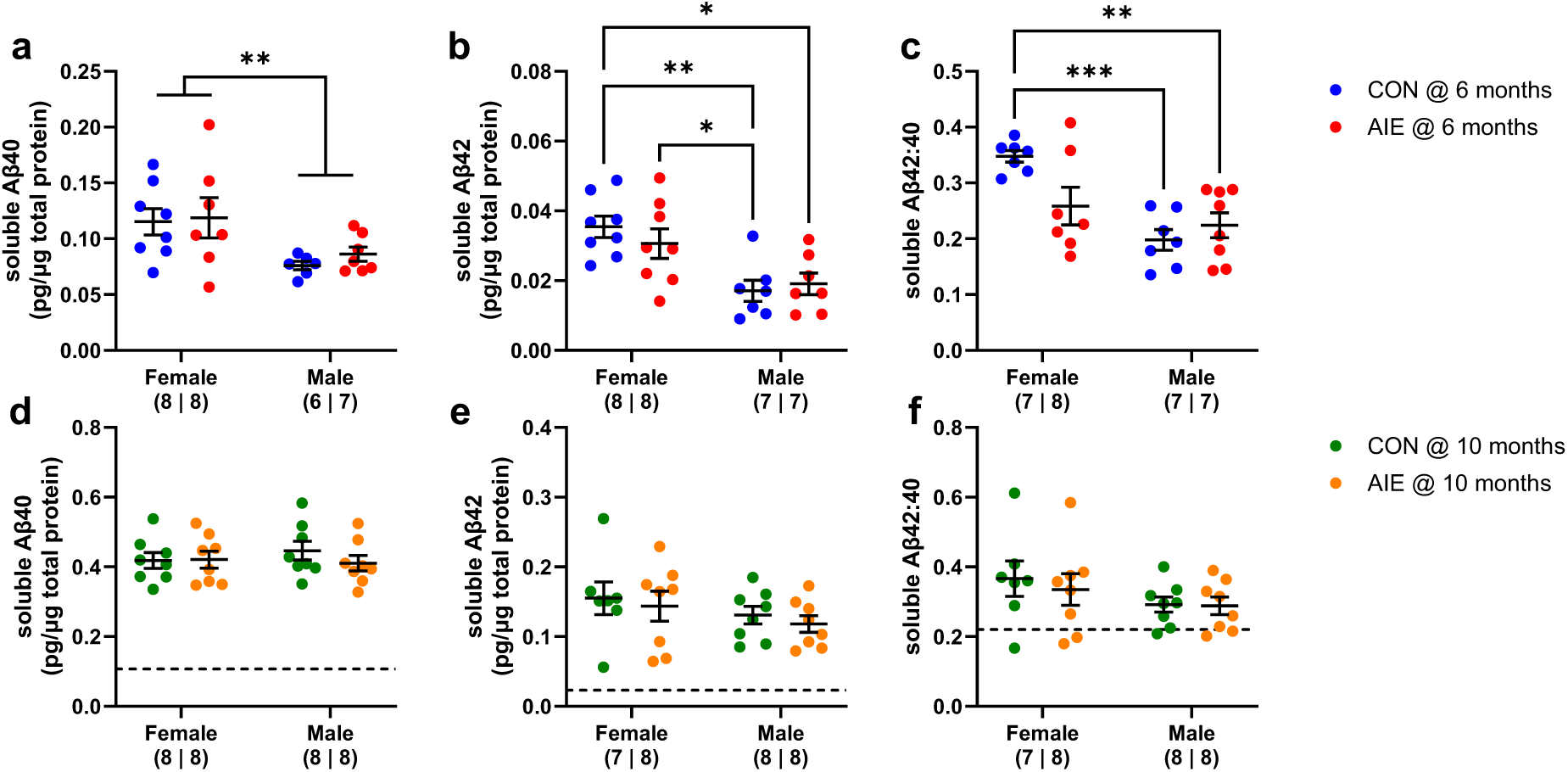
In the ventral hippocampus (vHPC), 6-month-old female TgF344-AD rats showed higher soluble Aβ40 and Aβ42 levels than males. In 6-month-old TgF344-AD rats, females showed higher Aβ40 (a; 2-way ANOVA, Main effect of sex: p = 0.0061), Aβ42 (b; 2-way ANOVA, Main effect of sex: p = 0.0002) levels than males, and CON females had a higher Aβ42/40 ratio than CON- and AIE-treated males (c; Main effect of sex : p = 0.0005; Treatment x Sex interaction: p = 0.0186 ). In 10-month-old TgF344-AD rats there were no differences in Aβ40, Aβ42, or Aβ42/40 between groups (d-f). There were age-dependent increases in Aβ40 (3-way ANOVA, Main effect of age: p = 0.0001), Aβ42 (3-way ANOVA, Main effect of age: p < 0.0001), and Aβ42/40 (3-way ANOVA, Main effect of age: p = 0.0030). Dashed line represents mean values of 6-month-old rats. ***p<0.001, **p < 0.01

However, in 10-month-old rats, there were no differences between groups in vHPC levels of Aβ40 (Figure 6d) or Aβ42 (Figure 6e), nor in the Aβ42/40 ratio (Figure 6f). While the early-life sex differences we observed at 6 months were no longer detectable in 10-month-old TgF344-AD rats, a 3-way ANOVA revealed a similar age-dependent increase in Aβ40 (Main effect of Age: F(1,53) = 541.5859, p = 0.0001), Aβ42 (Main effect of Age: F(1,53) = 146.1296, p < 0.0001), and Aβ42/40 (Main effect of age: F(1,52) = 9.7288, p = 0.0030;).

Lastly, in the dHPC, 6-month-old TgF344-AD female rats showed greater Aβ40 (Figure 7a; Main effect of Sex: F (1, 27) = 18.15; p = 0.0002;) and Aβ42 levels (Figure 7b; Main effect of Sex: F(1, 27) = 8.637; p = 0.0067) than TgF344-AD male rats. There were no differences between groups in the dHPC Aβ42/40 ratio (Figure 7c). Conversely, in 10-month-old TgF344-AD rats, AIE-treated rats had higher dHPC Aβ40 levels than control animals (Figure 7d; Main effect of Sex: F(1,27) = 6.751; p = 0.0150; Treatment x Sex interaction: F (1, 27) = 7.382; p = 0.0114). Post-hoc tests revealed that AIE-treated females had higher Aβ40 levels than control females (p = 0.0138), control males (p = 0.0167), and AIE-treated males (p = 0.0010). AIE-treated females also showed the highest dHPC Aβ42 levels (Figure 7e; Main effect of Sex: F(1,27) = 5.065; p = 0.0328; Treatment x Sex interaction: F (1, 27) = 15.15; p = 0.0006) and post-hoc tests revealed that AIE-treated females had higher Aβ42 levels than control females (LSD, p = 0.0074) and AIE-treated males (p = 0.0012). Despite this, there were no differences between groups in the Aβ42/40 ratio. Similar to the previous regions, a 3-way ANOVA revealed age-dependent increases in Aβ40 (Main effect of Age: F(1,57) = 4.7423, p = 0.0336), Aβ42 (Main effect of Age: F(1,57) = 13.0739, p = 0.0006;), and Aβ42/40 (Main effect of Age: F(1,57) = 25.1753, p < 0.0001). Collectively these studies demonstrate that, in TgF344-AD females, a history of AIE treatment leads to exacerbated Aβ40 and Aβ42 levels with age, with the dHPC being the most vulnerable region. This is the first study to identify the dHPC as an especially vulnerable region to alcohol-associated changes in AD-related pathology.

**Figure 7:**
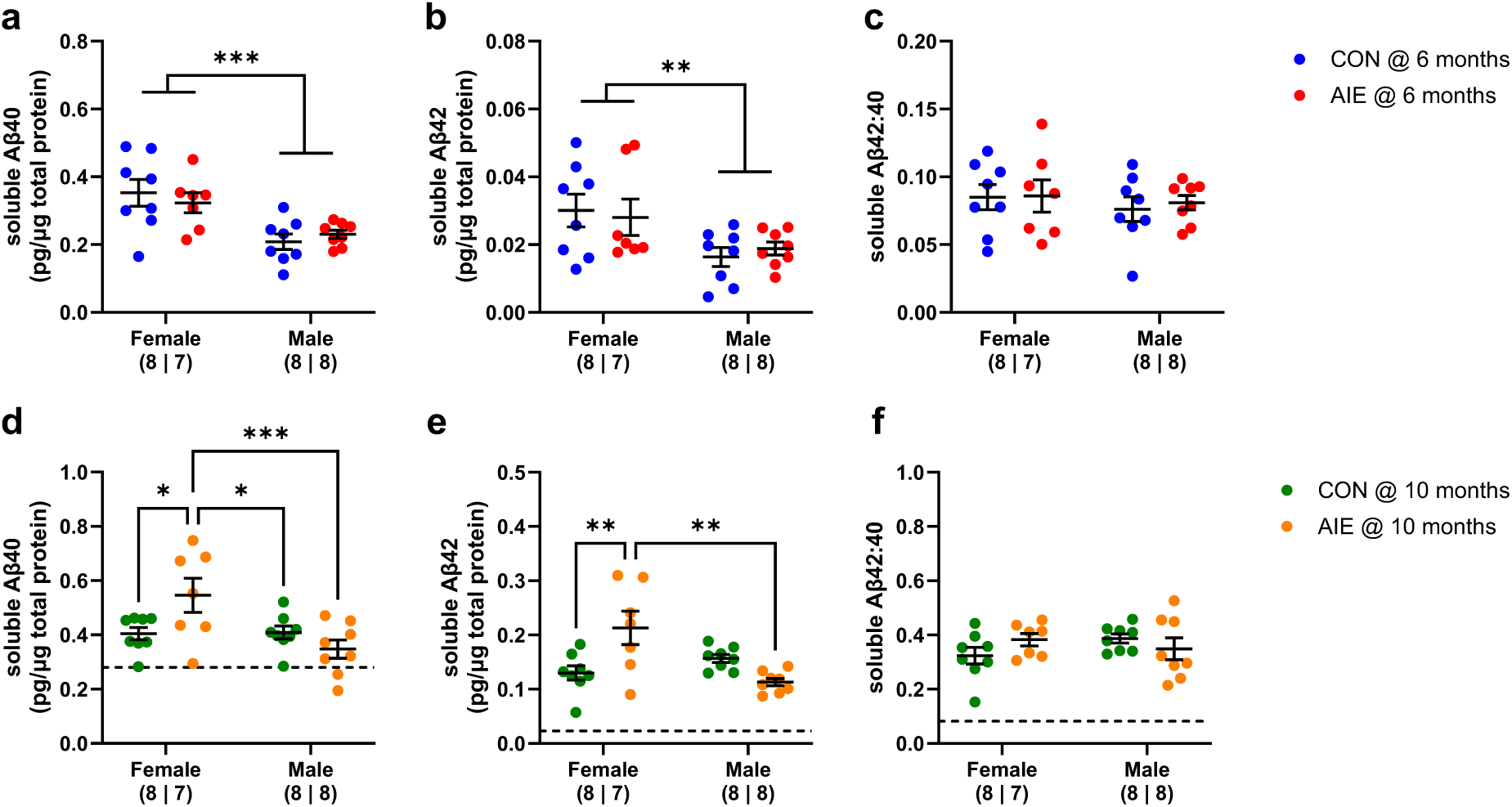
In the dorsal hippocampus (dHPC), 6-month-old female TgF344-AD rats showed higher soluble Aβ40 and Aβ42 levels than males. In 6-month-old TgF344-AD rats, females showed higher Aβ40 (a; 2-way ANOVA, Main effect of sex: p = 0.0002) and Aβ42 (b; 2-way ANOVA, Main effect of sex: p = 0.0067) levels compared to males. There were no differences between groups in Aβ42/40 at 6 months (c). In 10-month-old TgF344-AD rats Aβ40 levels were higher in AIE-treated females than CON-treated females, CON-treated males, and AIE-treated males (d; 2-way ANOVA, Main effect of sex: p = 0.015; Treatment x Sex: p = 0.0114), and Aβ42 levels were higher in AIE-treated females than CON-treated females and AIE-treated males (e; 2-way ANOVA, Main effect of sex: p = 0.0328; Treatment x Sex: p = 0.0006). However, there were no overall differences between groups in the Aβ42/40 ratio at 10 months (f). There were age-dependent increases in Aβ40 (3-way ANOVA, Main effect of age: p = 0.0336), Aβ42 (3-way ANOVA, Main effect of age: p = 0.0006), and Aβ42/40 (3-way ANOVA, Main effect of age: p < 0.0001). Dashed line represents mean values of 6-month-old rats. ***p<0.001, **p < 0.01, *p < 0.05

### Experiment 2: ISF Aβ40 levels change in AIE-treated rats after an acute ethanol exposure

Given that the dHPC is especially vulnerable to AIE-induced changes in AD-related pathology, we next used in vivo microdialysis to measure how ISF Aβ40 levels change in the dHPC after an acute ethanol exposure in control and AIE-treated females. Previous studies revealed that acute ethanol selectively increases ISF Aβ40 and not ISF Aβ42 levels [13]. Therefore, we limited this study to ISF Aβ40 levels. After an ethanol challenge (2.0 g/kg EtOH, i.p) AIE-treated rats showed a withdrawal-associated increase in ISF Aβ40 levels. In AIE-treated TgF334-AD rats, ISF Aβ40 levels increased by ∼157% from baseline at 4.5 hours post-EtOH, and remained elevated at ∼167% from baseline at 9.0 hours post-EtOH (Figure 8). A 3-way ANOVA revealed a Main Effect of Time (F (8, 144) = 4.600, p < 0.0001) and a Time x EtOH-challenge interaction (F (8, 144) = 2.270, p = 0.0257). Post-hoc tests revealed ISF that Aβ40 levels in the AIE+2.0 g/kg EtOH group were higher than CON + Saline group beginning at 4.5 hours post-EtOH; AIE+2.0 g/kg EtOH group were higher than AIE + Saline group beginning at 6.0 hours post-EtOH; and AIE+2.0 g/kg EtOH group were higher than CON + 2.0 g/kg EtOH group beginning at 7.5 hours post-EtOH (see supplementary table 1). We previously observed that acute ethanol leads to a withdrawal-induced increase in ISF Aβ40 levels, before returning to baseline [13]. Here, we observed a similar increase in ISF Aβ40 levels that corresponds to ethanol withdrawal; However, the protracted elevation in ISF Aβ40 levels might suggest that AIE disrupts Aβ clearance mechanisms in the dorsal hippocampus.

**Figure 8:**
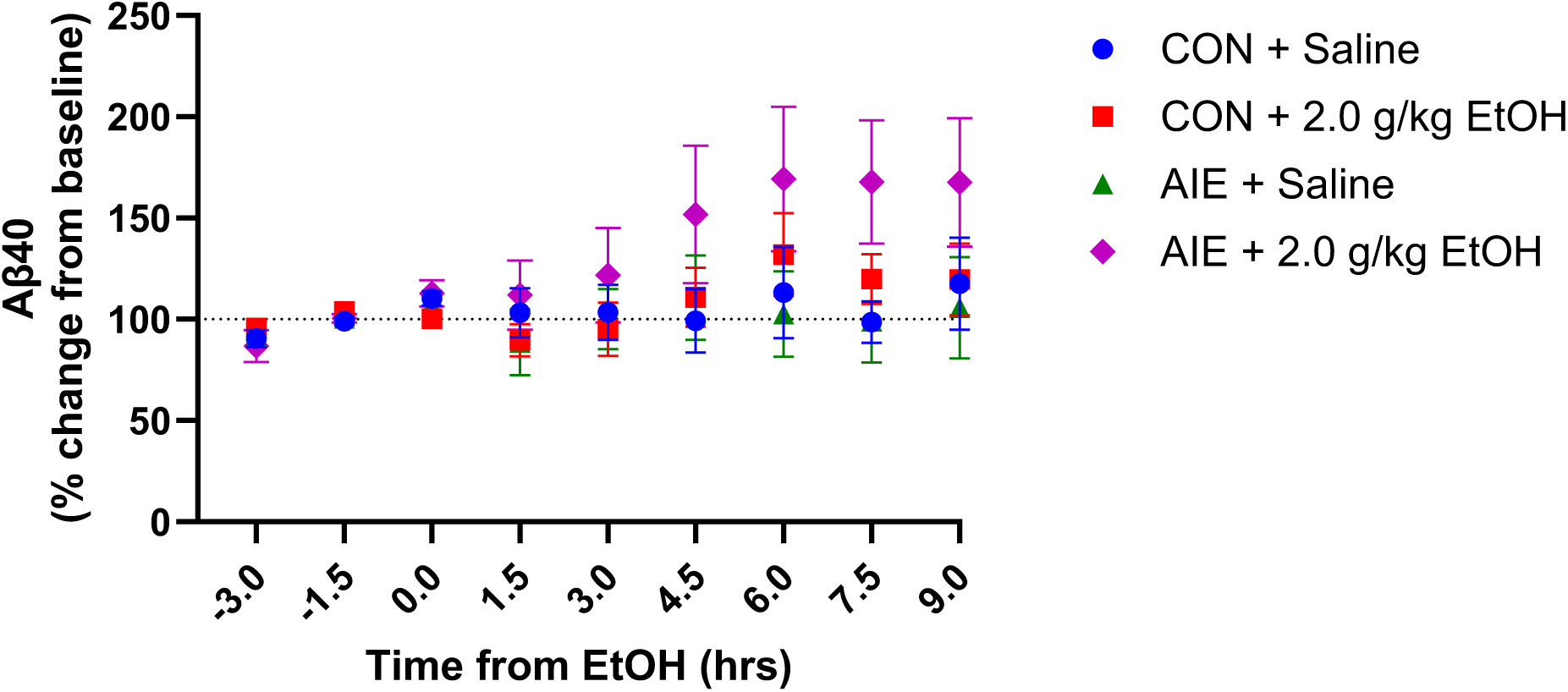
AIE-treated rats show dose-dependent increases in ISF Aβ40 levels after an acute ethanol exposure. In 3-month-old female Tg344-AD rats an acute ethanol challenge (EtOH-challenge), does not impact ISF Aβ40 levels in CON-treated females but did increase ISF Aβ40 levels in AIE-treated females over a 9 hour period (3-way ANOVA, Main effect of time: p < 0.0001; Time x EtOH-challenge: p = 0.0257). Fisher’s LSD showed that AIE-treated rats receiving 2.0 g/kg EtOH showed higher ISF Aβ40 levels than all other groups at 7.5- and 9.0-hours post-EtOH.

## Discussion

This study demonstrates that binge-like ethanol consumption during adolescence has age-, sex-, and brain region-dependent effects on Aβ levels in TgF344-AD rats. First, this study demonstrated that soluble Aβ levels are higher in female TgF344-AD rats at an early age in all regions tested, except in the mPFC. In female and male TgF344-AD rats, soluble Aβ levels increased with age throughout the brain - most notably in the PC. In the dHPC, the early sex differences in Aβ40 and Aβ42 were exacerbated with age in AIE-treated female rats (Figure 7). Thus, this study identifies the dHPC in female TgF344-AD rats as an especially sensitive region to AIE-dependent increases in Aβ. Lastly, this study demonstrated that an acute ethanol exposure induces a sustained increase in ISF Aβ40 levels in AIE-treated female TgF344-AD rats in the dHPC (Figure 8). To our knowledge, this is the first study to show the vulnerability of the dHPC to increases in Aβ in AIE-treated female Tg344-AD rats. These findings also indicate that the AIE-associated vulnerability may begin with impaired Aβ clearance early in life. These findings build on existing studies to demonstrate that binge-like ethanol consumption during adolescence can have long-term consequences on AD-related pathology.

In humans, women are at higher risk for AD, and show greater pathology and a more rapid advancement of clinical symptoms than men [20–22]. Those findings are mirrored in transgenic mouse models of Aβ overexpression, which show age- and sex-dependent increases in Aβ levels and amyloid burden in the hippocampus [39]. This study supports those findings in that we show female TgF344-AD rats have greater Aβ levels than TgF344-AD males at 6 months in the OFC, PC, EC and vHPC. By 10 months, these differences were no longer detectable in the OFC, PC, EC or vHPC, likely due to the age-dependent increase in Aβ in this model [25]. Interestingly, there were no differences in Aβ levels between sex or treatment groups in the mPFC. Here, Aβ40 and Aβ42 levels were highest in the dHPC of AIE-treated female TgF344-AD rats. Thus, this study identifies the dHPC as a particularly vulnerable region to alcohol-induced increases in AD-related pathology.

In AD patients, the hippocampus is vulnerable to amyloid deposition and neurodegeneration, with the dHPC being particularly sensitive to neurodegeneration [40]. Preclinical studies show that, as a whole, the hippocampus is also vulnerable to ethanol-associated increases in Aβ [13,19,41]. This study builds on these studies by showing that there are differences in ethanol-induced changes in Aβ levels between the dHPC and vHPC. At 6 months, which represents the prodromal period of AD pathogenesis in this model, female TgF344-Ad rats show higher Aβ40 and Aβ42 levels in both dHPC and vHPC, irrespective of AIE history (Figure 6a-c; 7a-c). By 10 months, however, these differences were no longer detectable in the vHPC while there was a clear Treatment x Sex interaction in the dHPC (Figure 6d-f; 7d-f). These data indicate that there may be differences in vulnerability between the dHPC and vHPC to alcohol-associated alterations in Aβ levels.

In recent years, several studies have begun to examine the differences in sensitivity between the dHPC and vHPC in rodents. Acute withdrawal (24-hours) from a 10-day chronic intermittent ethanol vapor exposure has sex- and region-specific effects on excitability in the dHPC and vHPC. In the vHPC, acute withdrawal results in a reduction in excitatory neurotransmission in female rats, while leading to an increase in excitatory neurotransmission in male rats. Meanwhile, in the dHPC acute withdrawal has no effect on excitatory in female rate, while decreasing excitability in male rats [42,43]. Given that these studies reveal sex- and region-specific effects on excitability in the dHPC and vHPC, it is conceivable that there may be further changes in excitability during a protracted withdrawal period. Thus, it is important to further investigate sex differences in ethanol-associated changes in excitability between the dHPC and vHPC.

Because we observed that AIE treatment exclusively increased Aβ levels in the dHPC of female TgF344-AD rats, we next used in vivo microdialysis to test whether acute ethanol had any impact on ISF Aβ levels in AIE-treated females early in life. Acute ethanol did not evoke any changes in ISF Aβ40 levels in ethanol-naïve female TgF344-AD rats. This discrepancy may be due to a number of factors including differences in sex (male vs female) or species (mouse vs rat). Future studies should seek to understand why acute ethanol evokes an increase in ISF Aβ levels in ethanol-naïve male mice, but not ethanol-naïve female rats. On the other hand, AIE-treated female TgF344-AD rats showed a dose-dependent increase in evoked ISF Aβ40 levels. Like the mouse studies, acute ethanol evoked an increase in ISF Aβ40 levels during acute ethanol withdrawal (3.0 – 6.0 hours post-ethanol). Unlike the mouse studies, ISF Aβ40 levels did not return to baseline but remained elevated for the remainder of the experiment (6.0 – 9.0 hours post-ethanol).

In AIE-treated rats, this response to acute ethanol reveals a potential mechanism by which AIE exacerbates Aβ levels in TgF344-AD female rats. First, acute ethanol exposure leads to a withdrawal-induced increase in ISF Aβ levels. These findings echo previous work demonstrating a transient ethanol withdrawal-associated increase in ISF Aβ40 levels [13]. Broadly speaking, ethanol bidirectionally modulates neuronal activity during intoxication and withdrawal by GABA-mediated inhibition during intoxication and increasing NMDA-mediated neuronal activity during withdrawal [44–50]. Given that Aβ40 is released from the brain in an activity-dependent manner, acute withdrawal in AIE-treated rats may be driving the activity-dependent increase in ISF Aβ40.

In ethanol-naïve APP/PS1 mice the withdrawal-associated increase in ISF Aβ40 was transient and returned to baseline levels within a few hours [13]. Whereas in this study, ISF Aβ40 levels increased during acute withdrawal and remained elevated for the remainder of the experiment. Here, we hypothesize that AIE may promote Aβ deposition by disrupting Aβ clearance mechanisms. Aβ is cleared from the brain through a number of active and passive mechanisms. These include extracellular degradation, microglial uptake and degradation, astrocytic uptake, astrocyte- and endothelial cell-mediated blood brain barrier transport, and passive bulk flow elimination through the CSF [51–53]. Thus, we hypothesize that binge-like ethanol consumption during adolescence exacerbates Aβ deposition via a two-hit model wherein AIE leaves the brain in a hyperexcitable state and impairs Aβ clearance mechanisms.

Preclinical studies have begun to examine the effects of ethanol on some of these Aβ clearance mechanisms, with the primary focus being on neuroinflammation as a driver of ethanol-associated Aβ aggregation. The impact of ethanol on neuroinflammation is well-known and has been reviewed extensively elsewhere [54–56]. Similarly, neuroinflammation is associated with increased Aβ deposition. In 7-month-old female 3xTg-AD mice, AIE treatment led to increased proinflammatory gene expression in the hippocampus that were positively correlated with Aβ levels [19]. In other preclinical studies a proinflammatory state promotes AD-related pathology, while increasing β- and γ-secretase activity and suppressing α-secretase activity [57]. Therefore, AIE treatment may be promoting Aβ production and inhibiting clearance by promoting neuroinflammation.

A few studies have begun exploring the impact of ethanol on other Aβ clearance mechanisms. In the brain, apolipoprotein E (ApoE) regulates amyloid deposition through a number of pathways and may inhibit Aβ clearance and promote the proliferation of amyloid plaques [58]. In preclinical studies, the impact of alcohol consumption or exposure on ApoE levels is mixed. In one study, ethanol consumption did not impact plasma ApoE levels in mice or humans with a history of alcohol misuse [59]. In another study, low-to-moderate ethanol vapor exposure over 3 days led to reduced ApoE levels in the cortex, and was associated with an overall reduction in Aβ level and plaque burden [60]. Despite this preliminary work, it is currently unknown how binge-like ethanol during adolescence impacts ApoE functionality in the context of AD-like pathology. Despite this growing body of research, the impact of chronic ethanol on the vast majority of Aβ clearance mechanisms remains largely unexplored and represents a significant gap in knowledge.

In support of previous findings [13–15], this study demonstrates that AIE leads to increased Aβ levels in 10-month-old female TgF344-AD rats. Importantly, this study identifies vulnerability differences in hippocampal subregions, with the dorsal hippocampus being most vulnerable to AIE-associated increases in Aβ. This sensitivity to Aβ may be apparent as early as 3-months, wherein an acute ethanol challenge led to sustained increases in Aβ in AIE-treated female TgF344-AD rats. Small perturbations in Aβ production and clearance can have an additive effect over a lifetime, therefore these studies suggest that the pathology may begin much earlier than expected. These findings contribute to a growing body of evidence demonstrating that alcohol misuse represents an important modifiable risk factor for AD. Future studies will focus on exploring the mechanisms by which binge-like ethanol consumption during adolescence directly influences Aβ production and clearance mechanisms.

## Acknowledgements

Amyloid-β (Aβ) antibodies were a generous gift from Hong Jiang and David Holtzman at Washington University in St. Louis.

## Author Contributions

Stephen M. Day (Conceptualization; Investigation; Formal analysis; Writing – original draft; Writing – review & editing; Funding acquisition); Nicole L. Reitz (Investigation; Writing – review & editing); Lisa M. Savage (Supervision; Conceptualization; Investigation; Formal analysis; Writing – original draft; Writing – review & editing; Funding acquisition).

## Ethical Considerations

The Binghamton University Institutional Animal Care and Use Committee approved the experimental procedures used in this study (approval no. 905-24) on May 27, 2024. All animal housing and experiments were conducted in strict accordance with the institutional Guidelines for Care and Use of Laboratory Animals at Binghamton University.

## Declaration of Conflicting Interests

The authors declared no potential conflicts of interest with respect to the research, authorship, or publication of this article.

## Funding

We would like to acknowledge the following funding sources: U01AA028710-04S1 (SMD, LMS), P50AA017823 (SMD), U01AA028710 (LMS).

## Data Availability Statement

The data supporting the findings of this study are available on request from the corresponding author.

---

**Supplementary Figure 1:**
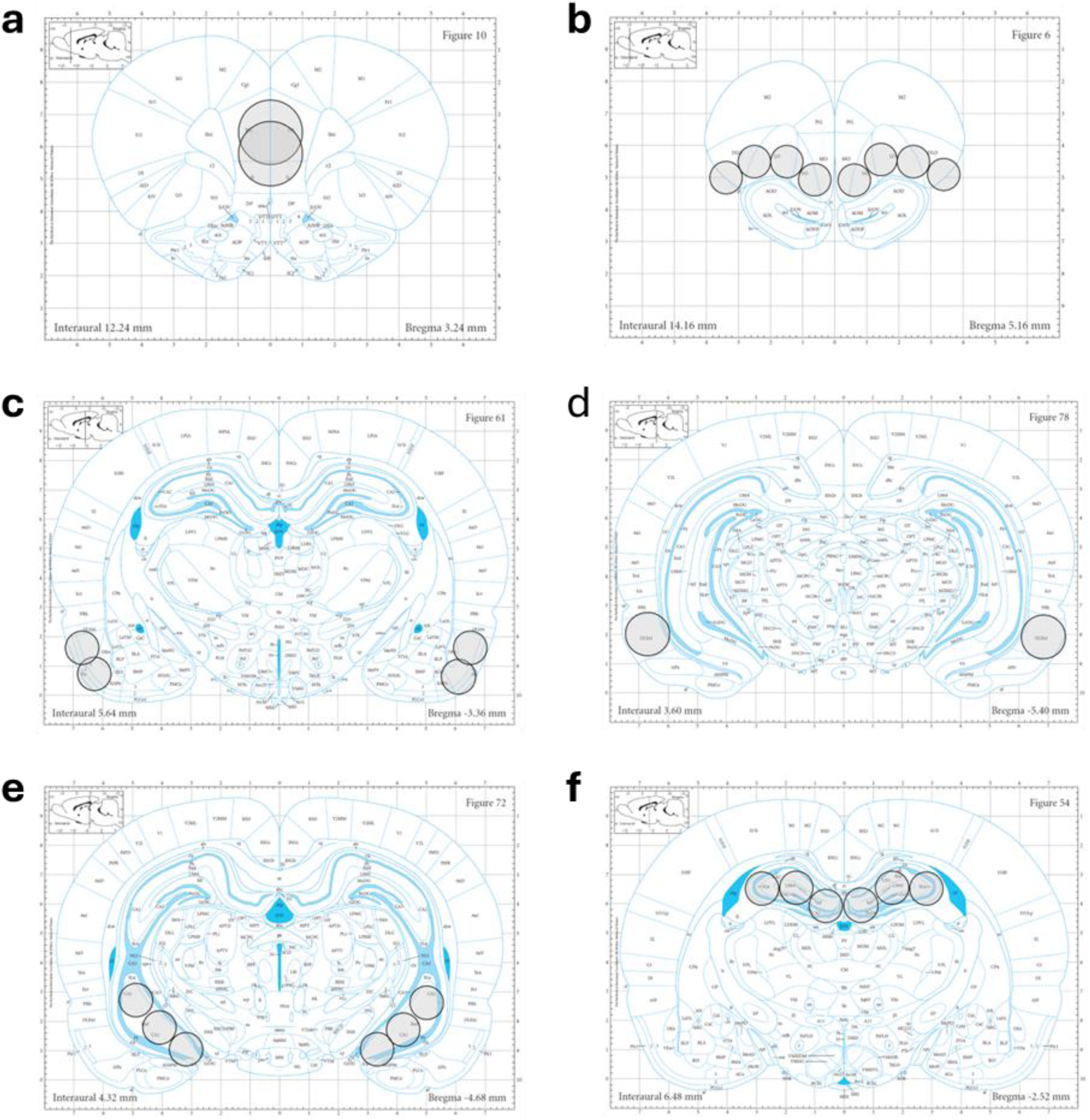
Regions selected for tissue punching determined by Paxinos & Watson 2013. Regions targeted for tissue punches determined by Paxinos & Watson 2013 [35]. a) Medial prefrontal cortex (Bregma, 3.24 mm); b) Orbitofrontal cortex (Bregma, 5.16 mm); c) piriform cortex (Bregma, -3.36 mm); d) entorhinal cortex (Bregma, -5.40 mm) e) ventral hippocampus (Bregma, -4.68 mm); f) dorsal hippocampus (Bregma, -2.52 mm).

**Supplementary Table 1:**
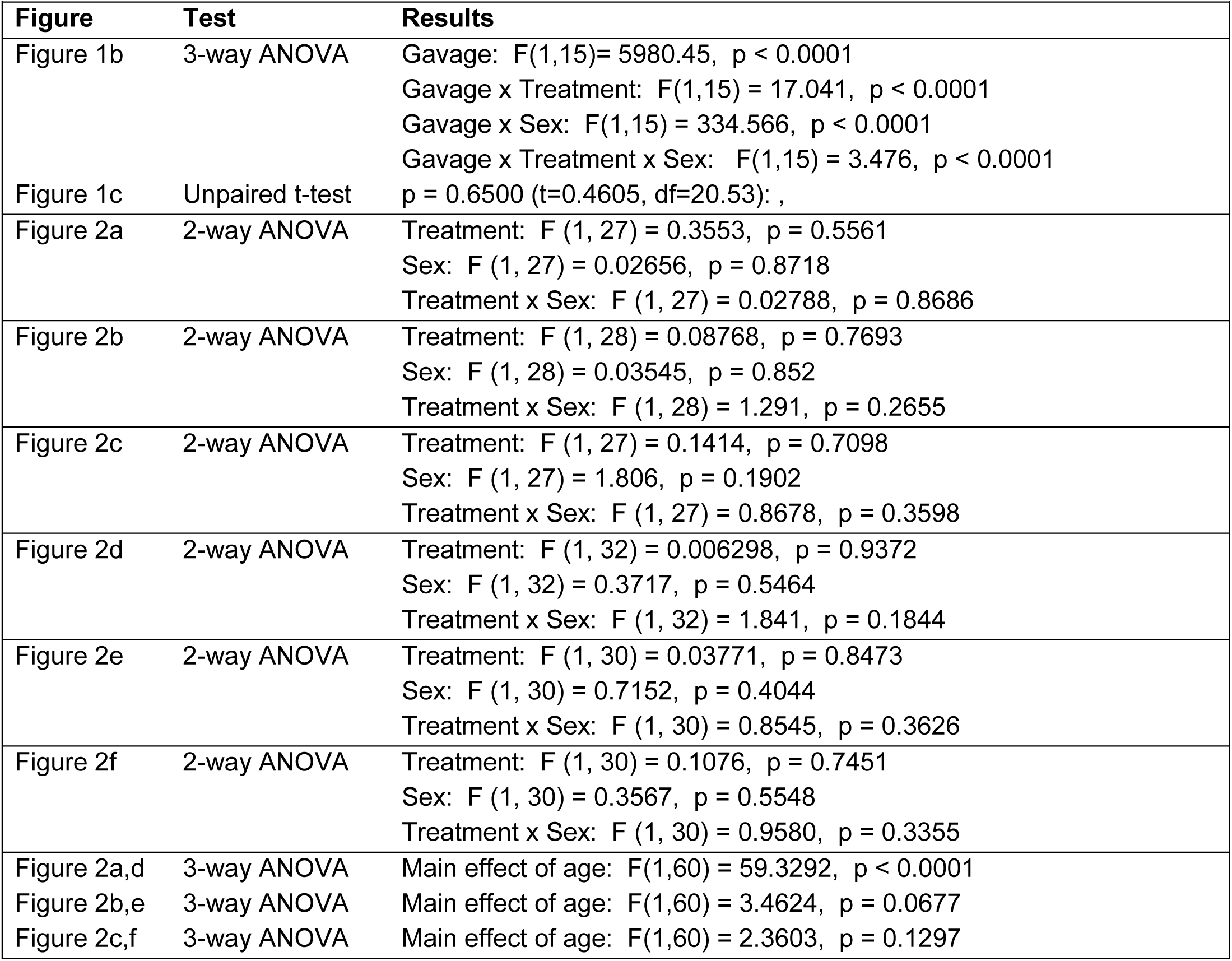
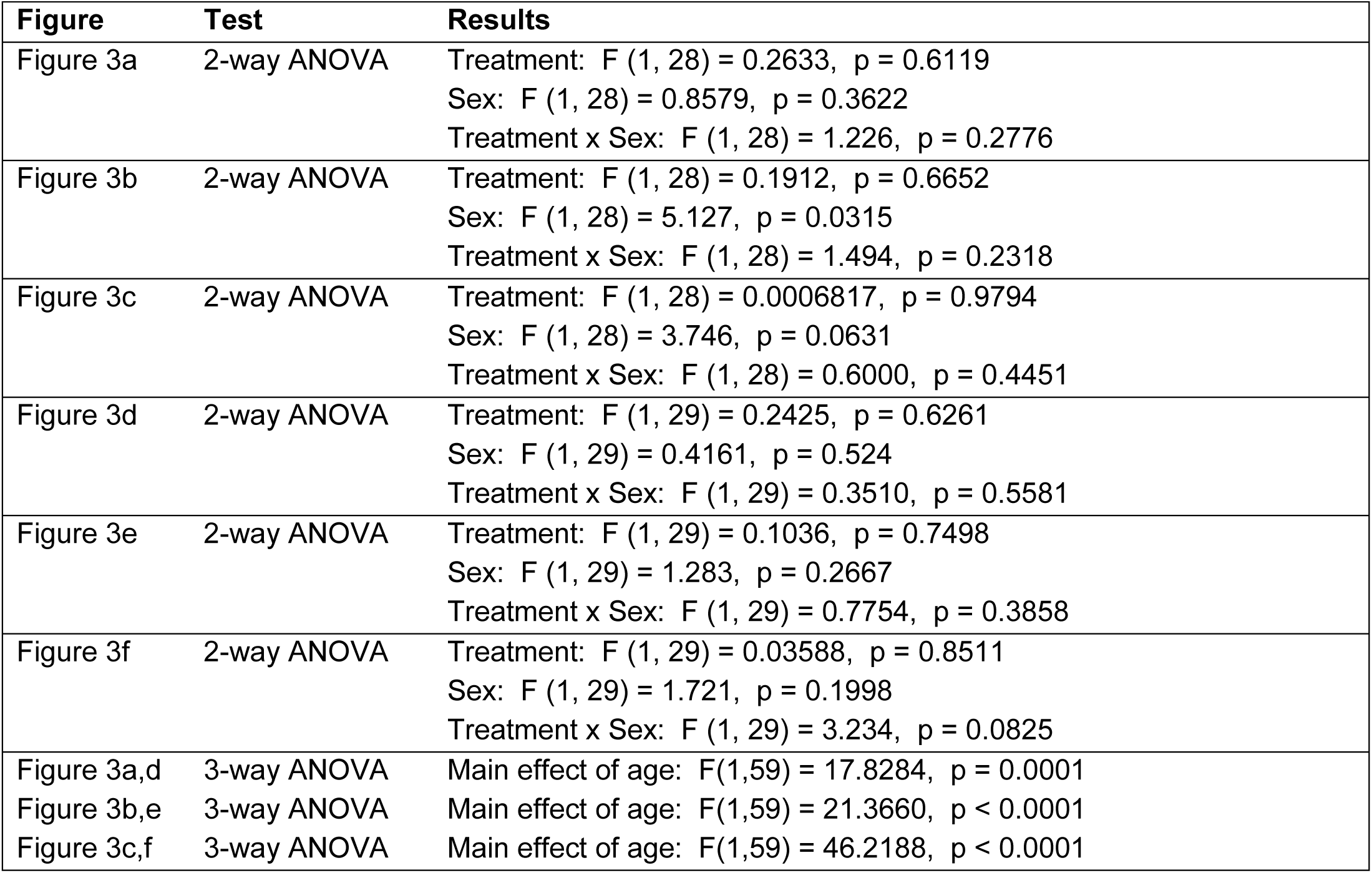
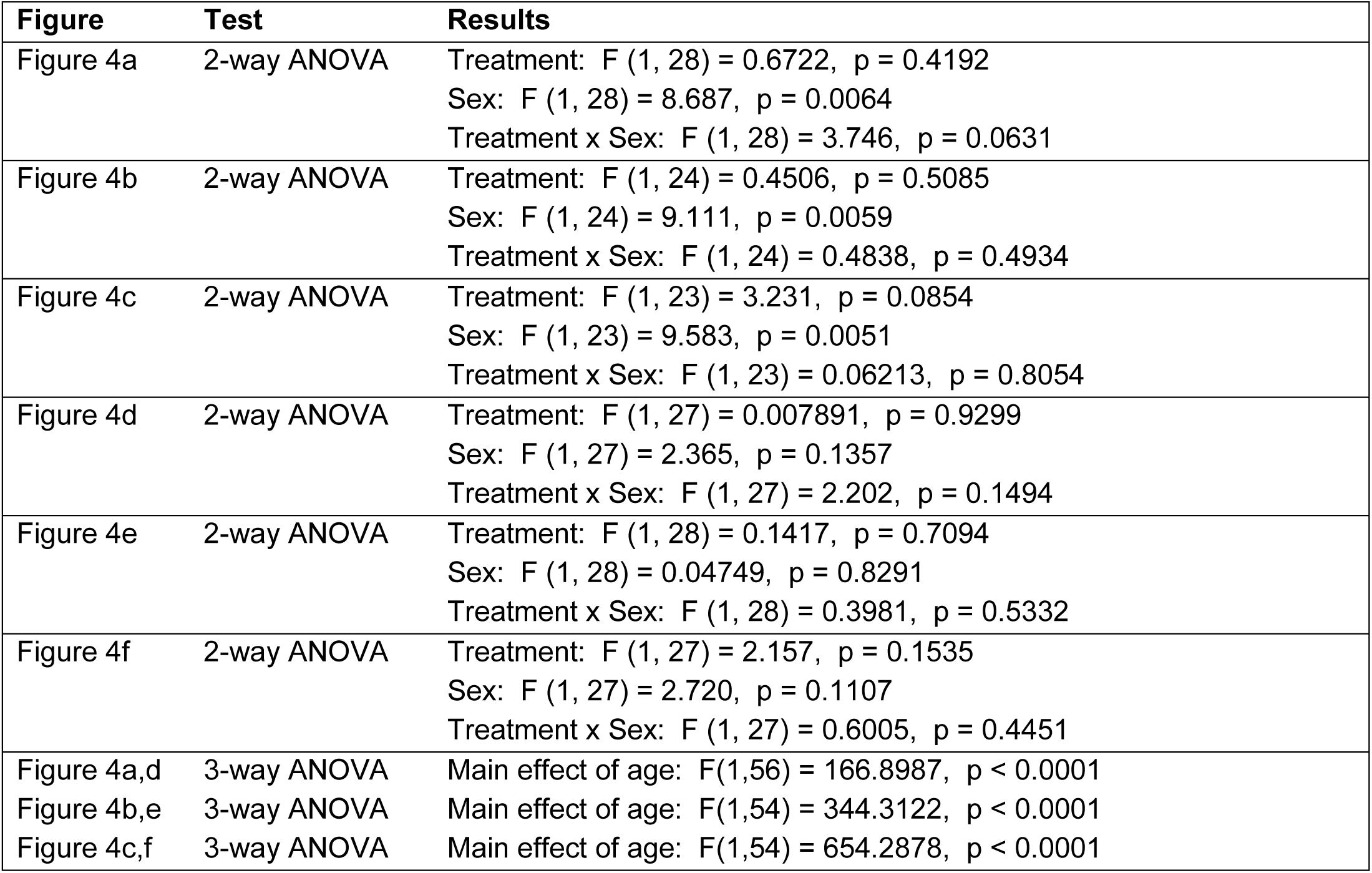
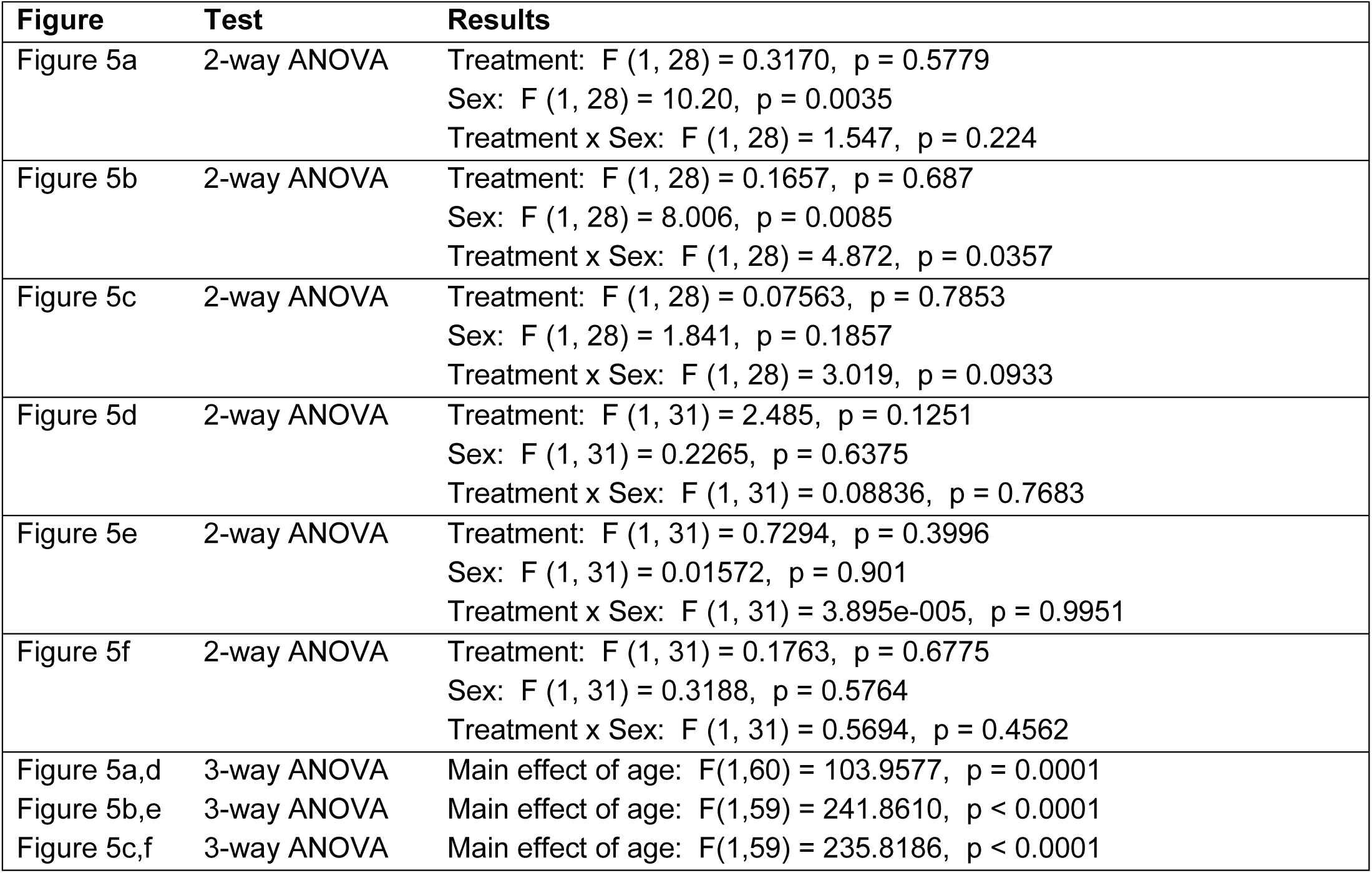
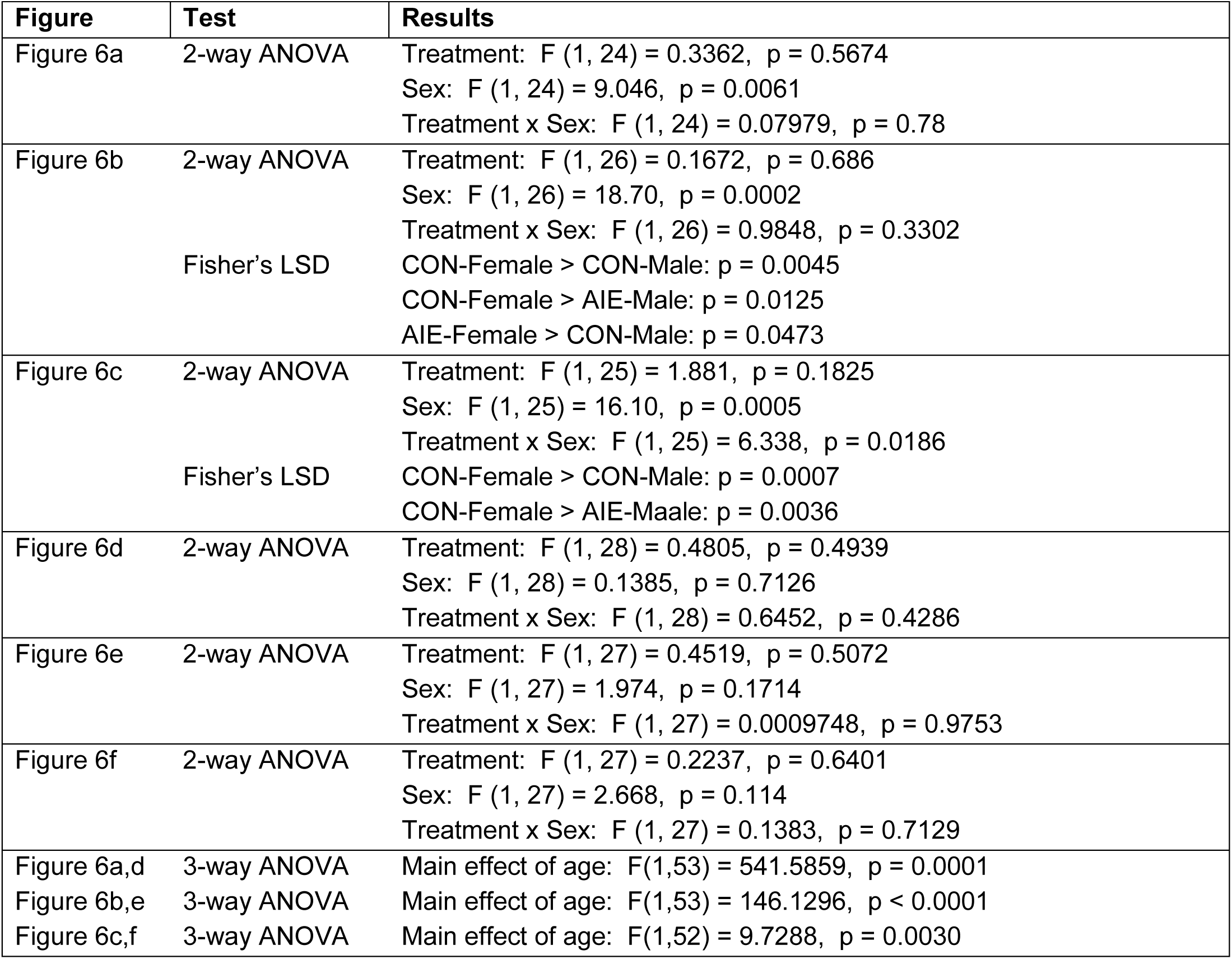
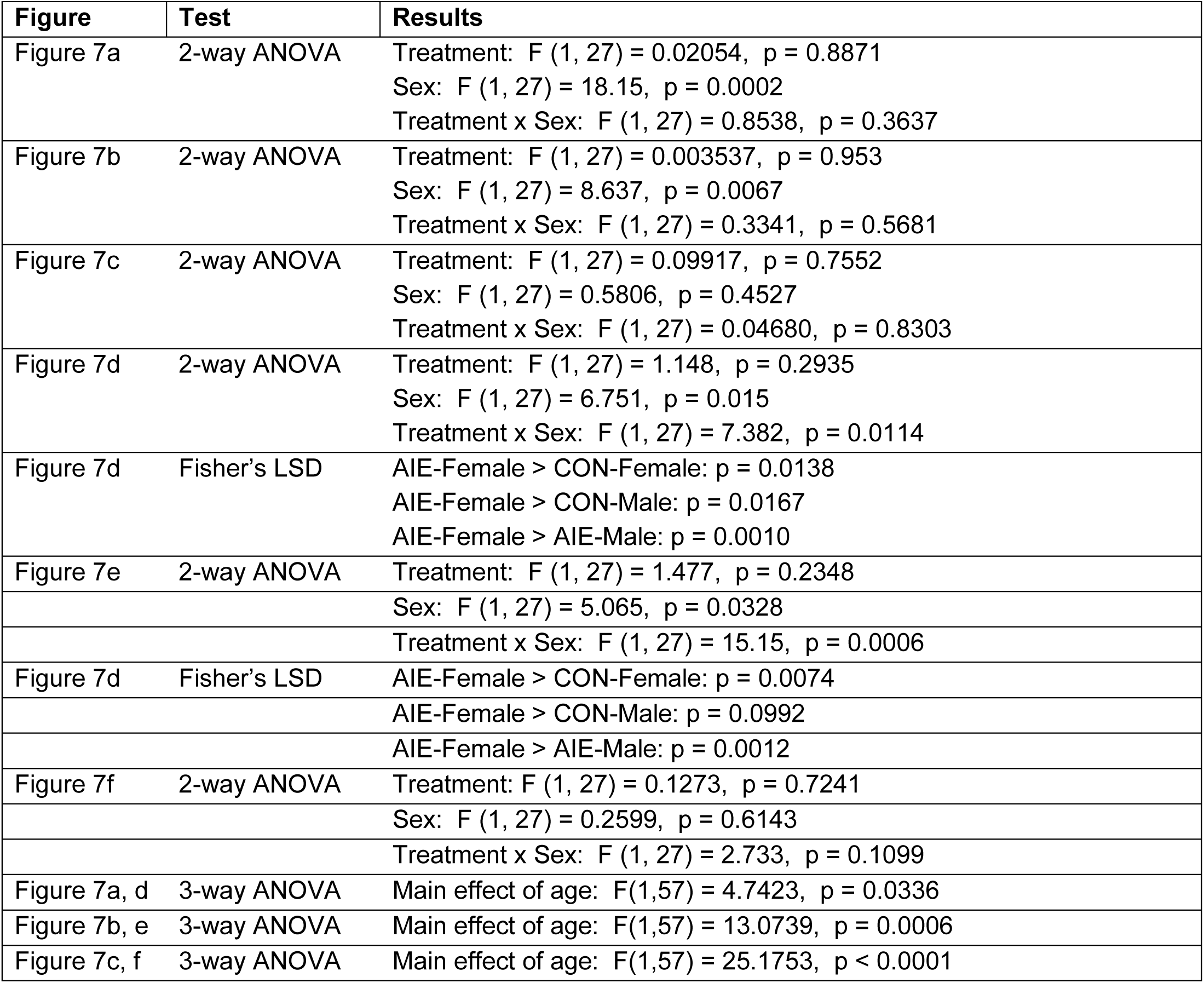
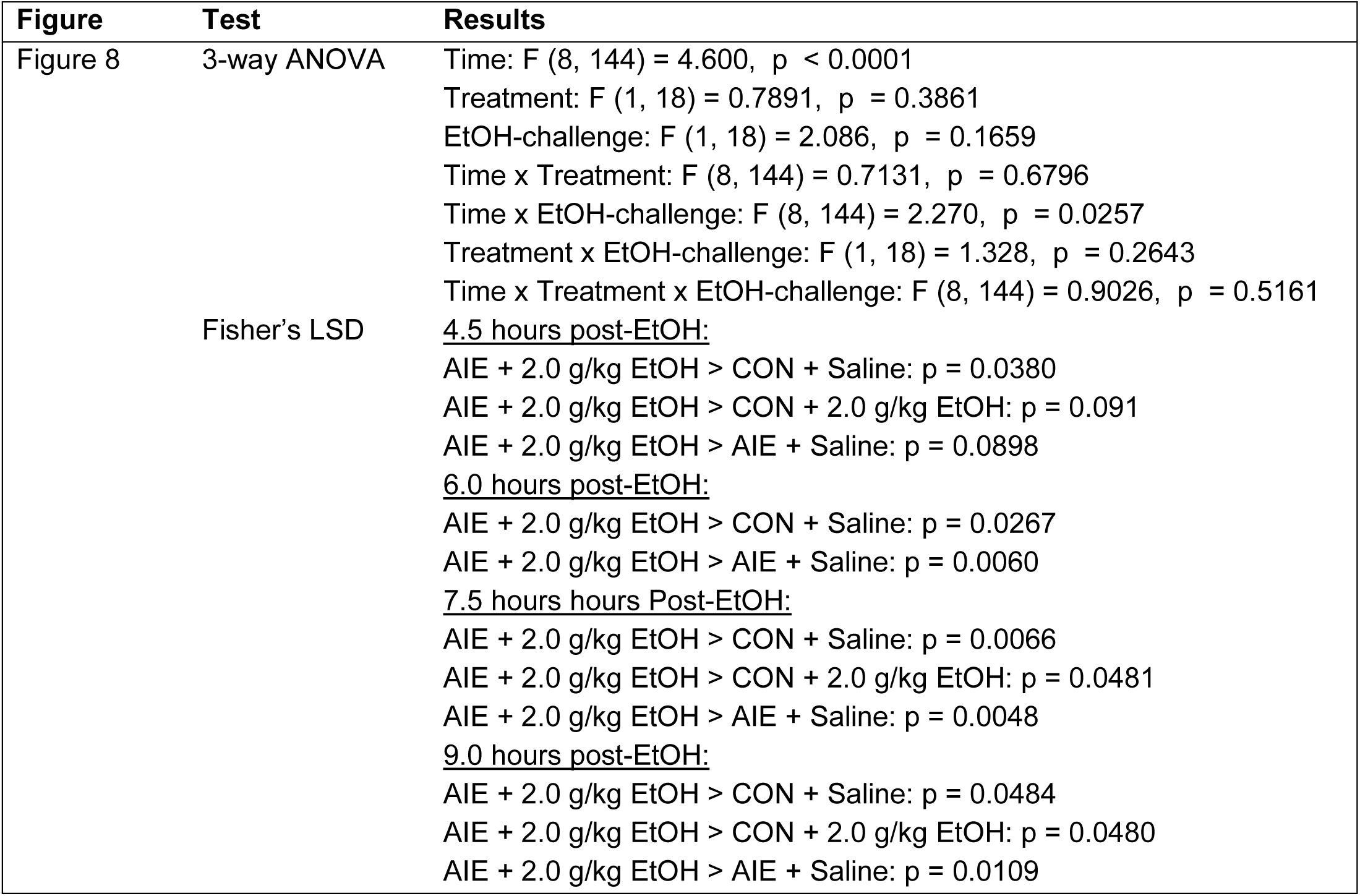
Statistical Results.

| **Supplementary Table 1: Statistical Results** |  |  |
| --- | --- | --- |
| **Figure** | **Test** | **Results** |
| Figure 1b | 3-way ANOVA | Gavage: $F(1,15) = 5980.45$, $p < 0.0001$<br>Gavage x Treatment: $F(1,15) = 17.041$, $p < 0.0001$<br>Gavage x Sex: $F(1,15) = 334.566$, $p < 0.0001$<br>Gavage x Treatment x Sex: $F(1,15) = 3.476$, $p < 0.0001$ |
| Figure 1c | Unpaired t-test | $p = 0.6500$ ($t=0.4605$, $df=20.53$): , |
| Figure 2a | 2-way ANOVA | Treatment: $F(1, 27) = 0.3553$, $p = 0.5561$<br>Sex: $F(1, 27) = 0.02656$, $p = 0.8718$<br>Treatment x Sex: $F(1, 27) = 0.02788$, $p = 0.8686$ |
| Figure 2b | 2-way ANOVA | Treatment: $F(1, 28) = 0.08768$, $p = 0.7693$<br>Sex: $F(1, 28) = 0.03545$, $p = 0.852$<br>Treatment x Sex: $F(1, 28) = 1.291$, $p = 0.2655$ |
| Figure 2c | 2-way ANOVA | Treatment: $F(1, 27) = 0.1414$, $p = 0.7098$<br>Sex: $F(1, 27) = 1.806$, $p = 0.1902$<br>Treatment x Sex: $F(1, 27) = 0.8678$, $p = 0.3598$ |
| Figure 2d | 2-way ANOVA | Treatment: $F(1, 32) = 0.006298$, $p = 0.9372$<br>Sex: $F(1, 32) = 0.3717$, $p = 0.5464$<br>Treatment x Sex: $F(1, 32) = 1.841$, $p = 0.1844$ |
| Figure 2e | 2-way ANOVA | Treatment: $F(1, 30) = 0.03771$, $p = 0.8473$<br>Sex: $F(1, 30) = 0.7152$, $p = 0.4044$<br>Treatment x Sex: $F(1, 30) = 0.8545$, $p = 0.3626$ |
| Figure 2f | 2-way ANOVA | Treatment: $F(1, 30) = 0.1076$, $p = 0.7451$<br>Sex: $F(1, 30) = 0.3567$, $p = 0.5548$<br>Treatment x Sex: $F(1, 30) = 0.9580$, $p = 0.3355$ |
| Figure 2a,d | 3-way ANOVA | Main effect of age: $F(1,60) = 59.3292$, $p < 0.0001$ |
| Figure 2b,e | 3-way ANOVA | Main effect of age: $F(1,60) = 3.4624$, $p = 0.0677$ |
| Figure 2c,f | 3-way ANOVA | Main effect of age: $F(1,60) = 2.3603$, $p = 0.1297$ |

| **Supplementary Table 1 (continued): Statistical Results** |  |  |
| --- | --- | --- |
| **Figure** | **Test** | **Results** |
| Figure 3a | 2-way ANOVA | Treatment: $F(1, 28) = 0.2633$, $p = 0.6119$<br>Sex: $F(1, 28) = 0.8579$, $p = 0.3622$<br>Treatment x Sex: $F(1, 28) = 1.226$, $p = 0.2776$ |
| Figure 3b | 2-way ANOVA | Treatment: $F(1, 28) = 0.1912$, $p = 0.6652$<br>Sex: $F(1, 28) = 5.127$, $p = 0.0315$<br>Treatment x Sex: $F(1, 28) = 1.494$, $p = 0.2318$ |
| Figure 3c | 2-way ANOVA | Treatment: $F(1, 28) = 0.0006817$, $p = 0.9794$<br>Sex: $F(1, 28) = 3.746$, $p = 0.0631$<br>Treatment x Sex: $F(1, 28) = 0.6000$, $p = 0.4451$ |
| Figure 3d | 2-way ANOVA | Treatment: $F(1, 29) = 0.2425$, $p = 0.6261$<br>Sex: $F(1, 29) = 0.4161$, $p = 0.524$<br>Treatment x Sex: $F(1, 29) = 0.3510$, $p = 0.5581$ |
| Figure 3e | 2-way ANOVA | Treatment: $F(1, 29) = 0.1036$, $p = 0.7498$<br>Sex: $F(1, 29) = 1.283$, $p = 0.2667$<br>Treatment x Sex: $F(1, 29) = 0.7754$, $p = 0.3858$ |
| Figure 3f | 2-way ANOVA | Treatment: $F(1, 29) = 0.03588$, $p = 0.8511$<br>Sex: $F(1, 29) = 1.721$, $p = 0.1998$<br>Treatment x Sex: $F(1, 29) = 3.234$, $p = 0.0825$ |
| Figure 3a,d | 3-way ANOVA | Main effect of age: $F(1,59) = 17.8284$, $p = 0.0001$ |
| Figure 3b,e | 3-way ANOVA | Main effect of age: $F(1,59) = 21.3660$, $p < 0.0001$ |
| Figure 3c,f | 3-way ANOVA | Main effect of age: $F(1,59) = 46.2188$, $p < 0.0001$ |
| **Figure** | **Test** | **Results** |
| Figure 4a | 2-way ANOVA | Treatment: $F(1, 28) = 0.6722$, $p = 0.4192$<br>Sex: $F(1, 28) = 8.687$, $p = 0.0064$<br>Treatment x Sex: $F(1, 28) = 3.746$, $p = 0.0631$ |
| Figure 4b | 2-way ANOVA | Treatment: $F(1, 24) = 0.4506$, $p = 0.5085$<br>Sex: $F(1, 24) = 9.111$, $p = 0.0059$<br>Treatment x Sex: $F(1, 24) = 0.4838$, $p = 0.4934$ |
| Figure 4c | 2-way ANOVA | Treatment: $F(1, 23) = 3.231$, $p = 0.0854$<br>Sex: $F(1, 23) = 9.583$, $p = 0.0051$<br>Treatment x Sex: $F(1, 23) = 0.06213$, $p = 0.8054$ |
| Figure 4d | 2-way ANOVA | Treatment: $F(1, 27) = 0.007891$, $p = 0.9299$<br>Sex: $F(1, 27) = 2.365$, $p = 0.1357$<br>Treatment x Sex: $F(1, 27) = 2.202$, $p = 0.1494$ |
| Figure 4e | 2-way ANOVA | Treatment: $F(1, 28) = 0.1417$, $p = 0.7094$<br>Sex: $F(1, 28) = 0.04749$, $p = 0.8291$<br>Treatment x Sex: $F(1, 28) = 0.3981$, $p = 0.5332$ |
| Figure 4f | 2-way ANOVA | Treatment: $F(1, 27) = 2.157$, $p = 0.1535$<br>Sex: $F(1, 27) = 2.720$, $p = 0.1107$<br>Treatment x Sex: $F(1, 27) = 0.6005$, $p = 0.4451$ |
| Figure 4a,d | 3-way ANOVA | Main effect of age: $F(1,56) = 166.8987$, $p < 0.0001$ |
| Figure 4b,e | 3-way ANOVA | Main effect of age: $F(1,54) = 344.3122$, $p < 0.0001$ |
| Figure 4c,f | 3-way ANOVA | Main effect of age: $F(1,54) = 654.2878$, $p < 0.0001$ |
| **Figure** | **Test** | **Results** |
| Figure 5a | 2-way ANOVA | Treatment: $F(1, 28) = 0.3170$, $p = 0.5779$<br>Sex: $F(1, 28) = 10.20$, $p = 0.0035$<br>Treatment x Sex: $F(1, 28) = 1.547$, $p = 0.224$ |
| Figure 5b | 2-way ANOVA | Treatment: $F(1, 28) = 0.1657$, $p = 0.687$<br>Sex: $F(1, 28) = 8.006$, $p = 0.0085$<br>Treatment x Sex: $F(1, 28) = 4.872$, $p = 0.0357$ |
| Figure 5c | 2-way ANOVA | Treatment: $F(1, 28) = 0.07563$, $p = 0.7853$<br>Sex: $F(1, 28) = 1.841$, $p = 0.1857$<br>Treatment x Sex: $F(1, 28) = 3.019$, $p = 0.0933$ |
| Figure 5d | 2-way ANOVA | Treatment: $F(1, 31) = 2.485$, $p = 0.1251$<br>Sex: $F(1, 31) = 0.2265$, $p = 0.6375$<br>Treatment x Sex: $F(1, 31) = 0.08836$, $p = 0.7683$ |
| Figure 5e | 2-way ANOVA | Treatment: $F(1, 31) = 0.7294$, $p = 0.3996$<br>Sex: $F(1, 31) = 0.01572$, $p = 0.901$<br>Treatment x Sex: $F(1, 31) = 3.895e-005$, $p = 0.9951$ |
| Figure 5f | 2-way ANOVA | Treatment: $F(1, 31) = 0.1763$, $p = 0.6775$<br>Sex: $F(1, 31) = 0.3188$, $p = 0.5764$<br>Treatment x Sex: $F(1, 31) = 0.5694$, $p = 0.4562$ |
| Figure 5a,d | 3-way ANOVA | Main effect of age: $F(1,60) = 103.9577$, $p = 0.0001$ |
| Figure 5b,e | 3-way ANOVA | Main effect of age: $F(1,59) = 241.8610$, $p < 0.0001$ |
| Figure 5c,f | 3-way ANOVA | Main effect of age: $F(1,59) = 235.8186$, $p < 0.0001$ |
| **Figure** | **Test** | **Results** |
| Figure 6a | 2-way ANOVA | Treatment: $F(1, 24) = 0.3362$, $p = 0.5674$<br>Sex: $F(1, 24) = 9.046$, $p = 0.0061$<br>Treatment x Sex: $F(1, 24) = 0.07979$, $p = 0.78$ |
| Figure 6b | 2-way ANOVA<br>Fisher's LSD | Treatment: $F(1, 26) = 0.1672$, $p = 0.686$<br>Sex: $F(1, 26) = 18.70$, $p = 0.0002$<br>Treatment x Sex: $F(1, 26) = 0.9848$, $p = 0.3302$<br>CON-Female > CON-Male: $p = 0.0045$<br>CON-Female > AIE-Male: $p = 0.0125$<br>AIE-Female > CON-Male: $p = 0.0473$ |
| Figure 6c | 2-way ANOVA<br>Fisher's LSD | Treatment: $F(1, 25) = 1.881$, $p = 0.1825$<br>Sex: $F(1, 25) = 16.10$, $p = 0.0005$<br>Treatment x Sex: $F(1, 25) = 6.338$, $p = 0.0186$<br>CON-Female > CON-Male: $p = 0.0007$<br>CON-Female > AIE-Male: $p = 0.0036$ |
| Figure 6d | 2-way ANOVA | Treatment: $F(1, 28) = 0.4805$, $p = 0.4939$<br>Sex: $F(1, 28) = 0.1385$, $p = 0.7126$<br>Treatment x Sex: $F(1, 28) = 0.6452$, $p = 0.4286$ |
| Figure 6e | 2-way ANOVA | Treatment: $F(1, 27) = 0.4519$, $p = 0.5072$<br>Sex: $F(1, 27) = 1.974$, $p = 0.1714$<br>Treatment x Sex: $F(1, 27) = 0.0009748$, $p = 0.9753$ |
| Figure 6f | 2-way ANOVA | Treatment: $F(1, 27) = 0.2237$, $p = 0.6401$<br>Sex: $F(1, 27) = 2.668$, $p = 0.114$<br>Treatment x Sex: $F(1, 27) = 0.1383$, $p = 0.7129$ |
| Figure 6a,d | 3-way ANOVA | Main effect of age: $F(1,53) = 541.5859$, $p = 0.0001$ |
| Figure 6b,e | 3-way ANOVA | Main effect of age: $F(1,53) = 146.1296$, $p < 0.0001$ |
| Figure 6c,f | 3-way ANOVA | Main effect of age: $F(1,52) = 9.7288$, $p = 0.0030$ |
| **Figure** | **Test** | **Results** |
| Figure 7a | 2-way ANOVA | Treatment: $F(1, 27) = 0.02054$, $p = 0.8871$<br>Sex: $F(1, 27) = 18.15$, $p = 0.0002$<br>Treatment x Sex: $F(1, 27) = 0.8538$, $p = 0.3637$ |
| Figure 7b | 2-way ANOVA | Treatment: $F(1, 27) = 0.003537$, $p = 0.953$<br>Sex: $F(1, 27) = 8.637$, $p = 0.0067$<br>Treatment x Sex: $F(1, 27) = 0.3341$, $p = 0.5681$ |
| Figure 7c | 2-way ANOVA | Treatment: $F(1, 27) = 0.09917$, $p = 0.7552$<br>Sex: $F(1, 27) = 0.5806$, $p = 0.4527$<br>Treatment x Sex: $F(1, 27) = 0.04680$, $p = 0.8303$ |
| Figure 7d | 2-way ANOVA | Treatment: $F(1, 27) = 1.148$, $p = 0.2935$<br>Sex: $F(1, 27) = 6.751$, $p = 0.015$<br>Treatment x Sex: $F(1, 27) = 7.382$, $p = 0.0114$ |
| Figure 7d | Fisher's LSD | AIE-Female > CON-Female: $p = 0.0138$<br>AIE-Female > CON-Male: $p = 0.0167$<br>AIE-Female > AIE-Male: $p = 0.0010$ |
| Figure 7e | 2-way ANOVA | Treatment: $F(1, 27) = 1.477$, $p = 0.2348$<br>Sex: $F(1, 27) = 5.065$, $p = 0.0328$<br>Treatment x Sex: $F(1, 27) = 15.15$, $p = 0.0006$ |
| Figure 7d | Fisher's LSD | AIE-Female > CON-Female: $p = 0.0074$<br>AIE-Female > CON-Male: $p = 0.0992$<br>AIE-Female > AIE-Male: $p = 0.0012$ |
| Figure 7f | 2-way ANOVA | Treatment: $F(1, 27) = 0.1273$, $p = 0.7241$<br>Sex: $F(1, 27) = 0.2599$, $p = 0.6143$<br>Treatment x Sex: $F(1, 27) = 2.733$, $p = 0.1099$ |
| Figure 7a, d | 3-way ANOVA | Main effect of age: $F(1,57) = 4.7423$, $p = 0.0336$ |
| Figure 7b, e | 3-way ANOVA | Main effect of age: $F(1,57) = 13.0739$, $p = 0.0006$ |
| Figure 7c, f | 3-way ANOVA | Main effect of age: $F(1,57) = 25.1753$, $p < 0.0001$ |

| Supplementary Table 1 (continued): Statistical Results |  |  |
| --- | --- | --- |
| Figure | Test | Results |
| Figure 8 | 3-way ANOVA | Time: $F(8, 144) = 4.600, p < 0.0001$<br>Treatment: $F(1, 18) = 0.7891, p = 0.3861$<br>EtOH-challenge: $F(1, 18) = 2.086, p = 0.1659$<br>Time x Treatment: $F(8, 144) = 0.7131, p = 0.6796$<br>Time x EtOH-challenge: $F(8, 144) = 2.270, p = 0.0257$<br>Treatment x EtOH-challenge: $F(1, 18) = 1.328, p = 0.2643$<br>Time x Treatment x EtOH-challenge: $F(8, 144) = 0.9026, p = 0.5161$ |
| | Fisher's LSD | <u>4.5 hours post-EtOH:</u><br>AIE + 2.0 g/kg EtOH > CON + Saline: $p = 0.0380$<br>AIE + 2.0 g/kg EtOH > CON + 2.0 g/kg EtOH: $p = 0.091$<br>AIE + 2.0 g/kg EtOH > AIE + Saline: $p = 0.0898$<br><u>6.0 hours post-EtOH:</u><br>AIE + 2.0 g/kg EtOH > CON + Saline: $p = 0.0267$<br>AIE + 2.0 g/kg EtOH > AIE + Saline: $p = 0.0060$<br><u>7.5 hours hours Post-EtOH:</u><br>AIE + 2.0 g/kg EtOH > CON + Saline: $p = 0.0066$<br>AIE + 2.0 g/kg EtOH > CON + 2.0 g/kg EtOH: $p = 0.0481$<br>AIE + 2.0 g/kg EtOH > AIE + Saline: $p = 0.0048$<br><u>9.0 hours post-EtOH:</u><br>AIE + 2.0 g/kg EtOH > CON + Saline: $p = 0.0484$<br>AIE + 2.0 g/kg EtOH > CON + 2.0 g/kg EtOH: $p = 0.0480$<br>AIE + 2.0 g/kg EtOH > AIE + Saline: $p = 0.0109$ |

**Supplementary Table 2:** Mean ISF Aβ40 levels in CON- and AIE-treated rats (see figure 8)

| Supplementary Table 2: Mean ISF Aβ40 levels in CON- and AIE-treated rats (see figure 8) | | | | | | | | |
| --- | --- | --- | --- | --- | --- | --- | --- | --- |
|  | CON + Saline<br>(n = 5) |  | CON + 2.0 g/kg EtOH<br>(n = 6) |  | AIE + Saline<br>(n = 6) |  | AIE + 2.0 g/kg EtOH<br>(n = 5) |  |
| Time | Mean | ± SEM | Mean | ± SEM | Mean | ± SEM | Mean | ± SEM |
| -3 | 90.55 | 4.21 | 95.83 | 2.71 | 92.05 | 2.41 | 86.7 | 7.98 |
| -1.5 | 99.12 | 1.84 | 103.85 | 0.99 | 100.45 | 2.56 | 100.49 | 2.13 |
| 0 | 110.34 | 3.62 | 100.32 | 2.49 | 107.5 | 3.75 | 112.81 | 6.46 |
| 1.5 | 103.24 | 12.19 | 89.73 | 7.97 | 88.7 | 16.21 | 111.89 | 17.17 |
| 3 | 103.49 | 13.67 | 95.12 | 13.12 | 100.08 | 14.83 | 121.75 | 23.36 |
| 4.5 | 99.3 | 15.68 | 110.95 | 14.54 | 110.79 | 20.89 | 151.79 | 33.98 |
| 6 | 113.23 | 22.6 | 132.03 | 20.54 | 102.53 | 21.08 | 169.36 | 35.76 |
| 7.5 | 98.73 | 10.25 | 119.99 | 12.44 | 99.1 | 20.54 | 167.83 | 30.4 |
| 9 | 117.67 | 22.72 | 119.7 | 17.69 | 105.66 | 25.08 | 167.57 | 31.74 |

## Notes

### Competing Interest Statement

The authors have declared no competing interest.

